# Bringing large scale top-down metabolic modelling to eukaryotes with PhotoEukStein: a meta-model for unicellular phototrophs

**DOI:** 10.1101/2023.05.22.541783

**Authors:** Marie Burel, Antoine Régimbeau, Sirine Boucenna, Céline Orvain, Nadia Perchat, Alain Perret, Adrien Thurotte, Damien Eveillard, Eric Pelletier

## Abstract

Defining biological functional traits for unicellular organisms relies on comprehending the set and combination of the biochemical reactions their genomes encode for. This network of biochemical reactions defines the metabolic strategy organisms and communities use. Understanding the functional traits of unicellular organisms involves studying the combination of biochemical reactions encoded in their genomes. These reactions determine the metabolic strategy that organisms and communities use to grow in a specific environment. While prokaryotes have been extensively studied for their metabolic networks, eukaryotes have lagged behind due to the complexity of their genomes and the need for a better understanding of their metabolism. We have created *PhotoEukstein*, a meta-metabolic model for unicellular phototrophic eukaryotes. This meta-model enables quick and automated derivation of Genome-Scale Metabolic models directly from genomes. We have used it to analyse 533 environmental genomes and marine eukaryotic unicellular plankton transcriptomes. These models can help predict functional traits that cannot be purely deducted from taxonomic information or the listing of metabolic reactions encoded by the genome. They provide the opportunity to build connections with Earth System Models to pinpoint environmental parameters to capture specific functional traits.

## Introduction

Marine plankton are the dominant life form in the ocean, covering a broad diversity of organisms from viruses up to meter-size cnidarians via archaea, bacteria and single-celled eukaryotes, and have highly dynamic interactions. Together, these organisms play an active role in maintaining the Earth’s system, carrying out almost half of the net primary production on the planet ^1^ and exporting photosynthetically fixed carbon to the deep oceans ^2^. Yet, a large part remains elusive to in-depth laboratory investigations. While ocean ecosystems biology investigates how biotic and abiotic processes determine emergent properties of the ocean ecosystem as a whole, the only biological knowledge we have from a large part of plankton comes from environmental genomics data.

With the ability to generate a vast amount of sequencing data out of environmental samples at ever-decreasing costs and the improvement of bioinformatics methods to reconstruct high-quality genomes from metagenomic data, several hundreds or thousands of Metagenome-Assembled Genomes (MAGs) have been reconstructed for viruses, bacteria, archaea, and eukaryotes, covering a significant fraction of the biological diversity in several environments ^3–12^ . These environmental genomes greatly expand genomic and transcriptomic knowledge of cultured organisms ^13,14^. Furthermore, most of these genomes correspond to organisms without cultured representatives, although they represent species contributing to the global biomass and cycling of nutrients. For example, while green algae and protists represent a third of the total marine biomass ^15^, eukaryote genomes recovered from marine environments differ from reference sequences and can even describe putative new phylum ^8^. Omics-based approaches can hence significantly contribute to gaining knowledge about the biology of these uncultured organisms.

However, one cannot mechanistically understand these organisms and how they interact with their environment through the sole prism of omics data. Functional annotation and phenotypic characterisation are essential to allow us to gain further insight into "who is doing what" rather than answering the question of "who is here." In particular, systems biology approaches have been instrumental in acquiring a detailed stoichiometric representation of metabolic phenotypes via constraint-based reconstruction and analysis ^16^.

As biological features (traits/phenotypes) of organisms are primarily driven by their metabolic abilities ^1,17^, reconstructing metabolic networks from environmental -omics data provides a unique way to study the biology and ecology of these organisms and communities, as well as to draw a better picture of their influence on the environment ^18^. Indeed, metabolic networks are the cornerstone of Genome-Scale Metabolic Models (GSMs), which have demonstrated numerous applications in various field, such as biotechnology or synthetic biology ^19^. GSMs are constraint-based models that use -omic knowledge in metabolic networks and reformulate it into linear inequalities. GSMs regroup all the metabolic reactions encoded in a genome or transcriptome, and their intertwining (cf. Methods, section 1). We can use tools such as Flux-Balanced Analysis (FBA) or Flux Variation Analysis (FVA) ^20^ (see ^19^ for a review) to explore the solution space. This space encompasses all the potential solutions in the n-dimensions space defined by metabolic fluxes that satisfy the constraints imposed on each flux in the GSM) ^21^. By doing this, we can predict metabolic phenotypes ^22^ by optimising an objective function, typically schematising the organismal growth rate. Nevertheless, it is essential to note that GSMs do not consider various biological regulations that modulate enzymatic activities within a cell, such as regulation of gene expression or protein synthesis, post-translational modifications of proteins, or protein-protein interactions).

However, behind its benefit, reconstructing a metabolic network from -omics data analysis is a tedious task initially performed only for reference genomes, mobilising tedious laboratory experiments and metabolism experts for long periods and requiring expertise dedicated to a single genome ^23,24^ for a review). With every new genome sequenced, the traditional bottom-up approach requires these tedious and time-consuming tasks are to be repeated. Metabolic modelling for eukaryotes has primarily been restricted to well-studied model organisms (*Homo sapiens*, *Arabidopsis thaliana, Phaeodactylum tricornutum, Saccharomyces cerevisiae* being the most complete examples). Few efforts have been devoted to unicellular phototroph organisms, even though they represent half of Earth’s net primary production ^25^.

As a recent alternative, top-down semi-automated approaches deriving GSMs from a global reference pan-genomic collection of described reactions have been proposed ^26^. The curation of such a generic model is performed only once. It is then converted into ready-for-use organism-specific models while preserving all manual curation and relevant structural properties. Among the most efficient algorithm for metabolic modelling, both in terms of computational time and quality of resulting models, is CarveMe ^26,27^, which has been used in various studies (for examples ^28–32^) to derive prokaryotic GSMs. The EMBL-GEMs database (https://github.com/cdanielmachado/embl_gems) encompasses more than 5500 bacterial or archaeal simulation-ready GSMs.

Here, we report the development of PhotoEukStein, a reference-based metabolic meta-model for unicellular eukaryotic phototrophs, and its use with the CarveMe algorithm to reconstruct constraint-based metabolic models on a collection of 259 MAGs and 274 transcriptomes, plus the 16 references. The analysis of this resource revealed that, while there is a taxonomic imprinting of the repertoire of metabolic reactions across the 549 organisms considered, metabolic network topologies of resulting GSMs suggest that distantly related organisms can display similar metabolic phenotypes in a given environmental context. While a metabolic framework is particularly well-suited to formalise and analyse an organism’s ecological niche, deriving GSMs for unicellular (phototroph) eukaryotes pave the way to describe their metabolic phenotypes and functional traits and ecology.

## Results

### PhotoEukStein rationale and validation of reconstructed genome-scale metabolic models

PhotoEukStein combines 16 existing available metabolic models of marine unicellular eukaryote phototrophs species ranging from Rhodophytes (red algae), Chlorophytes (green algae), Streptophytes, Stramenopiles (brown algae), and Haptophytes to Cryptophytes; along with *Arabidopsis thaliana* and *Klebsormidium nitens* (see Supplementary Table S1). These reference organisms cover most of the described taxonomical diversity of phototrophs eukaryotes (Figure 1a and Figure 1b). Out of these 16, 4 results from experts-curated bottom-up annotation, namely *Chlamydomonas reinhardtii* iRC1080 ^33^, *Chlorella variabilis* ^34^, *Phaeodactylum tricornutum* iLB1031 ^35^ and *Thalassiosira pseudonana* ^36^. After merging these models as per the protocol described in ^24^, a manual curation phase was performed to make PhotoEukStein ready for constraint-based analyses (see Material and Methods for details). PhotoEukStein encompasses 5831 metabolites and 11229 reactions, 7599 of the which being associated with 20468 protein sequences from reference genomes. Two types of reactions are distinguished: 2067 boundary reactions (including 360 exchanges reactions) accounting for the transport of metabolites from or to the environment, and 9162 internal metabolic processes (see Supplementary Table S3).

**Figure 1:**
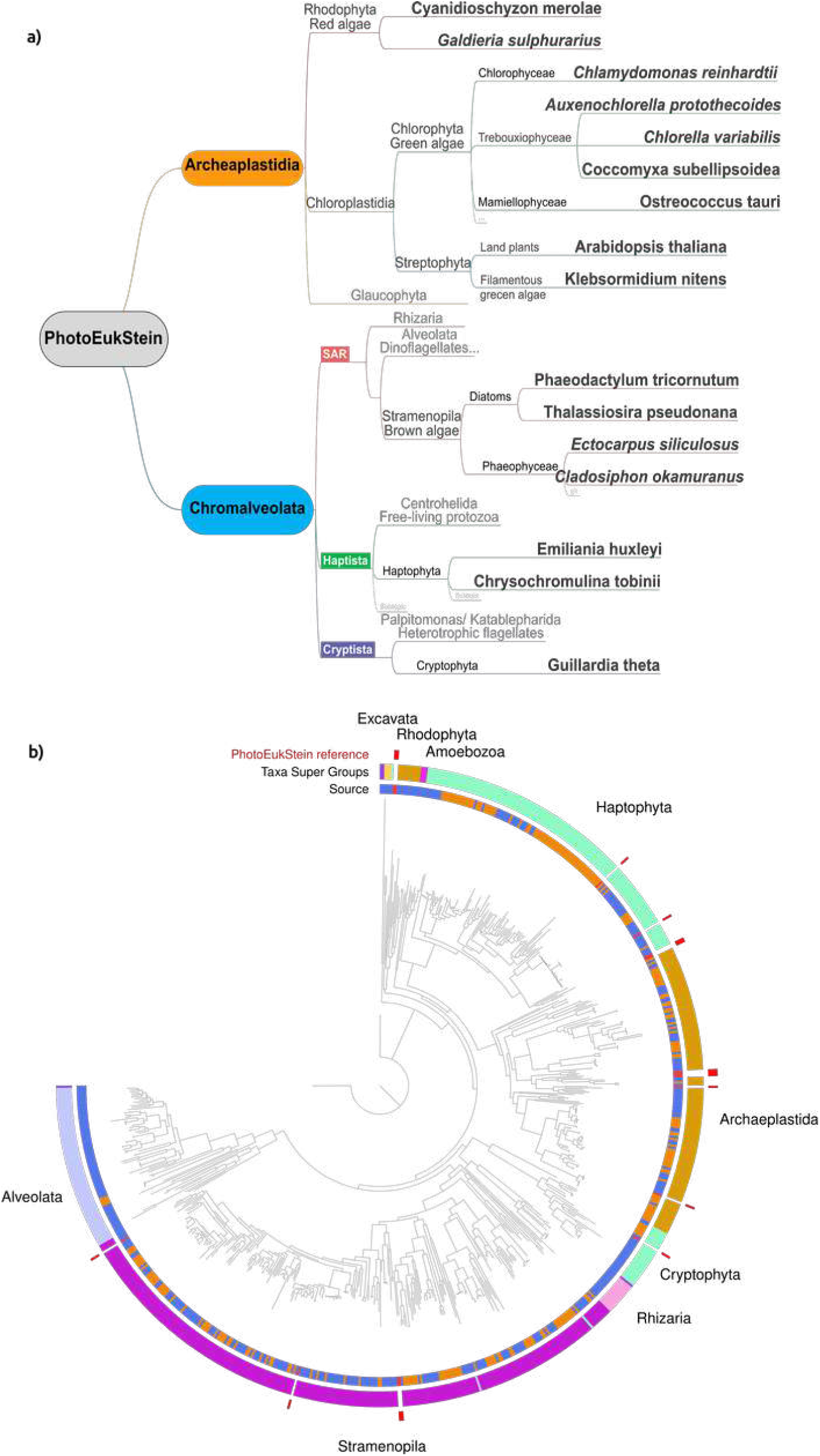
a) PhotoEukStein construction. Taxonomic diversity of the 16 existing GSMs that were combined to generate PhotoEukStein. b) Taxonomic diversity of the 549 PhotoeukStein-derived GSMs applied on 259 MAGs from Tara Oceans data (Delmont 2022, in orange in the inner circle) and 274 transcriptomes from METdb (Niang 2018, in blue in the inner circle). The taxonomic distribution of the 16 reference GSMs is indicated in ref (outer circle). Main taxonomic groups are indicated in the medium circle. Center is a dendrogram representing the taxonomy.

For all 16 model organisms used for the construction of PhotoEukStein, we derived GSM using the CarveMe algorithm. Nearly systematically (14 cases out of 16) more reactions and metabolites were included in PhotoEukStein-derived GSMs as compared to reference GSMs (from 1.5 to 6.7 times more reactions), a notable exception being *Guillardia theta* for which the reference GSM is only composed of 121 reactions (Supp. Table 2). Only two PhotoEukStein-based GSMs contain slightly fewer reactions than the reference one (*Cladosiphon okamuranus* - 86% and *Arabidopsis thaliana* - 98%). Interestingly, these two organisms are multicellular.

Several reasons can be invoked to explain this different number of considered reactions: i) ad-hoc expert-based models often focus on specific aspects of the metabolism (e.g., the lipids metabolism in *C. reinhardtii* ^33^), ii) specific choices made during PhotoEukStein construction phase, were, for example, reactions only appearing in the *A. thaliana* GSM were discarded from PhotoEukStein, as potentially representing specific terrestrial multicellular plants-specific reactions, and iii) only reactions with identified corresponding protein sequences encoded in the genome were considered during the expert-led curation of the reference GSMs. Indeed, the CarveMe process allowed the inclusion of the minimal set of reactions from the meta-model that are mandatory to maintain a functional GSM (i.e. with gap-less pathways), even if they lack the identification of associated proteins. Thus, the process may overcome incomplete gene predictions or partial MAG completions. Moreover, it can point to critical reactions with yet unidentified related genes in the genome.

For 3 out of the 16 reference GSMs (*Chlorella variabilis*, *Phaeodactylum tricornutum,* and *Thalassiosira pseudonana*), predicted growth rates were compared initially with culture-based data ^34–37^ (cf. Methods for details). To assess the validity of the PhotoEukStein-derived GSMs, we extensively sampled the metabolic niches and compared growth rates predicted from expert-based GSM with those obtained from PhotoEukStein-based GSMs (Supplementary Figure S1). Both predicted growth rates were highly correlated in all cases, showing that PhotoEukStein-derived GSMs are as accurate as expert-based models in capturing fundamental metabolic knowledge and *in vivo* observations from cultures. Therefore, PhotoEukStein-based GSMs are likely to provide valuable biological insights solely based on the genomic information of organisms.

Moreover, exploring metabolic niches with GSMs allows for assessing the metabolic exchange fluxes associated with higher growth rates, either for reference or PhotoEukStein-based GSMs. This delineate the environment’s metabolic limitations for organisms or missing metabolic reactions. For the three reference species we studied, metabolite exchange fluxes differentiating growth rates between both models vary, and a given metabolite can favour growth in either reference or PhotoEukStein GSM, depending on the organism.

To further scrutinise the internal consistency of PhotoEukStein-derived GSMs, we compared reaction fluxes as predicted by the manually curated (iLB1031) and automaticaly reconstructed models for *Phaeodactylum tricornutum* (for details, cf. Materials and Methods section 3). We considered fluxes correlations between reactions within each model by sampling the whole metabolic space (Supplementary Figure S2). The resulting correlation maps were highly similar in reference and PhotoEukStein-derived GSMs, indicating that reactions in both models are very similarly interconnected. We confirmed this observation by plotting intra-model correlations values distribution (Supplementary Figure S3). When pairs of reactions were highly correlated within one GSM, they were as highly correlated within the other GSM, and low connected pairs of reactions had similar characteristics in both models. The automated top-down approach applied to *P. tricornutum* captured the same essentials as the expert-based model. It represented the same biological features even when considering the distribution of metabolic fluxes within the GSM.

### Applying PhotoEukStein to new cultured species

Although *Chlorella variabilis*, *Phaeodactylum tricornutum,* and *Thalassiosira pseudonana* are three species whose experimentaly curated GSMs where included into data corpus used to build the PhotoEukStein meta-model, we investigated PhotoEukStein’s ability to predict growth rates for new unicellular algal species. *Pelagomonas calceolata* RCC100 (a pelagophyte) and *Teleaulax amphioxeia* AND-0710 (a cryptophyte) were selected precisely because they fall outside the scope of the models originally embedded in the tool, thus constituting an independent validation dataset. Using their complete genome sequences, we derived GSMs using PhotoEukStein for *P. calceolata* RCC100 and *T. amphioxeia* AND-0710. We explored their metabolic niches to predict the effects of nitrate and photon deprivation on cellular growth.

We grew cultures under varying light intensities and nitrogen (NO_3_) concentrations (cf. Materials and Methods section 4). We fitted logistic growth curves to the experimental data, yielding species-specific growth rate parameters (r) across the tested conditions (Supplementary Figures S4 and S5, respectively). Fit quality was consistently high across conditions for both species, as shown by the close overlap between used observed data points and fitted curves, despite minor discrepancies. Thus, minor deviations were observed at high light intensities for T. *amphioxeia* AND-0710, likely reflecting photo-inhibition or late-phase mortality not fully captured even after truncation. For *P. calceolata*, the condition L = 2, N = 0.1 yielded no detectable growth (r = NaN), consistent with severe co-limitation by light and nitrogen, and was excluded from subsequent analyses. Overall, the robustness of the logistic fits across 19-20 conditions per species supports the reliability of the extracted r values used for comparison with PhotoEukStein growth predictions.

Both species exhibited the expected dual dependency on light and nitrogen availability, with growth rates increasing with light intensity and peaking at intermediate nitrogen concentrations before declining at higher levels. We then compared these experimentally derived growth rate landscapes with the metabolic niche projections generated by PhotoEukStein for each species (Supplementary Figure S6). The model predictions, based on genome-scale metabolic reconstructions and fluxes of photon and nitrate uptake reactions, qualitatively recapitulate the observed growth patterns: both the experimental and predicted surfaces show low growth under light limitation, a steep increase in growth rate beyond a light threshold, and a region of maximal growth at high photon flux combined with moderate nitrate availability. The triangular feasible region in the model projections reflects the thermodynamic and stoichiometric constraints of the metabolic network and is consistent with the experimentally observed physiological envelope. Quantitative differences in absolute growth rate values between experiments and model predictions are expected, as genome-scale metabolic models optimise flux distributions rather than directly predict kinetic growth rates. Nevertheless, the qualitative concordance across two phylogenetically distinct species supports the applicability of the PhotoEukStein framework beyond its original species set.

### Reconstruction and analysis of 549 genome-scale metabolic models

Unicellular eukaryote phototrophs MAGs data from *Tara* Oceans were extracted from previous study ^8^, while unicellular eukaryotic phototrophs transcriptomes were recovered from the METdb database ^14^ which extends the MMETSP resource ^13^. In total, 259 MAGs and 274 transcriptomes, genomic data, along with the 16 organisms used as a reference to build PhotoEukStein (Supplementary Table S2), were used as input for the CarveMe method ^26^ to derive 549 dedicated GSMs from PhotoEukStein (for details, cf Materials and Method section 5).

The resulting GSMs contain a mean of 4154 reactions each (min. 1350, max. 7045), 72.7% of them (min. 41.6%, max 89.1%) being associated with a gene from the MAG/transcriptome input (as compared with the 67.67 % of PhotoEukStein reactions being associated with a reference sequence). As anticipated, the number of metabolic reactions in the resulting GSMs retained during the graph refinement (or carving) process decreased with the estimated level of genomes completion (Supplementary Table S2, Figure Extended Data). Nevertheless, when dealing with partially complete genomes, the intrinsic feature of CarveMe to keep reactions within the GSM even without supporting protein evidence allowed us to highlight mandatory reactions yet to be identified in a given genome.

The phylogeny mainly influences the repertoire of metabolic reactions across species as a given genome’s global repertoire of genes. Acquisition of new metabolic functions resulting from horizontal gene transfers from viruses ^38,39^ or bacteria ^40^, can superimpose alternative connectivity within the metabolic pathways. To assess the reliability of the functional capabilities of marine unicellular phototroph eukaryotes across taxonomy, we analysed the reaction content of PhotoEukStein-derived GSMs. First, we computed the Jaccard distance between our 549 GSMs based on the presence/absence of metabolic reactions (Supplementary Figure S7). We observe groups of reactions associated explicitly with some taxonomical groups (for example, reactions linked with lipid metabolism are specific to the diatoms). We sketch a more global picture of the distribution of metabolic reactions across our collection of GSMs, by performing a Uniform Manifold Approximation and Projection (UMAP) analysis ^41,42^ of the presence/absence of reactions within each GSM (as listed in Supplementary Table S3). This showed that GSMs are not evenly distributed in the functional space and that there is strong imprinting of the taxonomic origin of the corresponding organisms in the functional proximity (Supplementary Figure S8). This observation agrees with the vertical transmission of most metabolic-associated genes (for a review, see ^43^).

To further define the distribution of GSMs within the functional space, we performed a k-means clustering based on the presence/absence of reactions. Five clusters corresponding to metabolic groups were revealed (Supplementary Figure S8, Supplementary Table S2). When taxonomic information associated with each GSM is projected on the clusters, we observe a substantial taxonomic-based composition effect: most of the Stramenopiles are concentrated in cluster I (131 out of 212) and V (66 out of 212), Archaeplastida in clusters II (52 out of 118) and III (52 out of 118), Hacrobia in cluster IV (114 out of 135), and Alveolata and Amoebozoa in cluster V (respectively 40 out of 51 and 8 out of 10). Cluster V was the more taxonomically diverse, with eight taxonomic supergroups represented out of 10.

Beyond the presence/absence of reaction analysis of metabolic models, studying their topology provides insights into how metabolites are intertwined within the model. We explored the diversity of our 549 PhotoEukStein-derived GSMs using a diffusion map approach ^44,45^ to capture the non-linear combinations of metabolic capabilities variables represented in a 549-dimension space representing the internal metabolic reactions of GSMs. We then proceeded to a dimension reduction to visualise that diversity using the UMAP algorithm (Figure 3).

**Figure 2:**
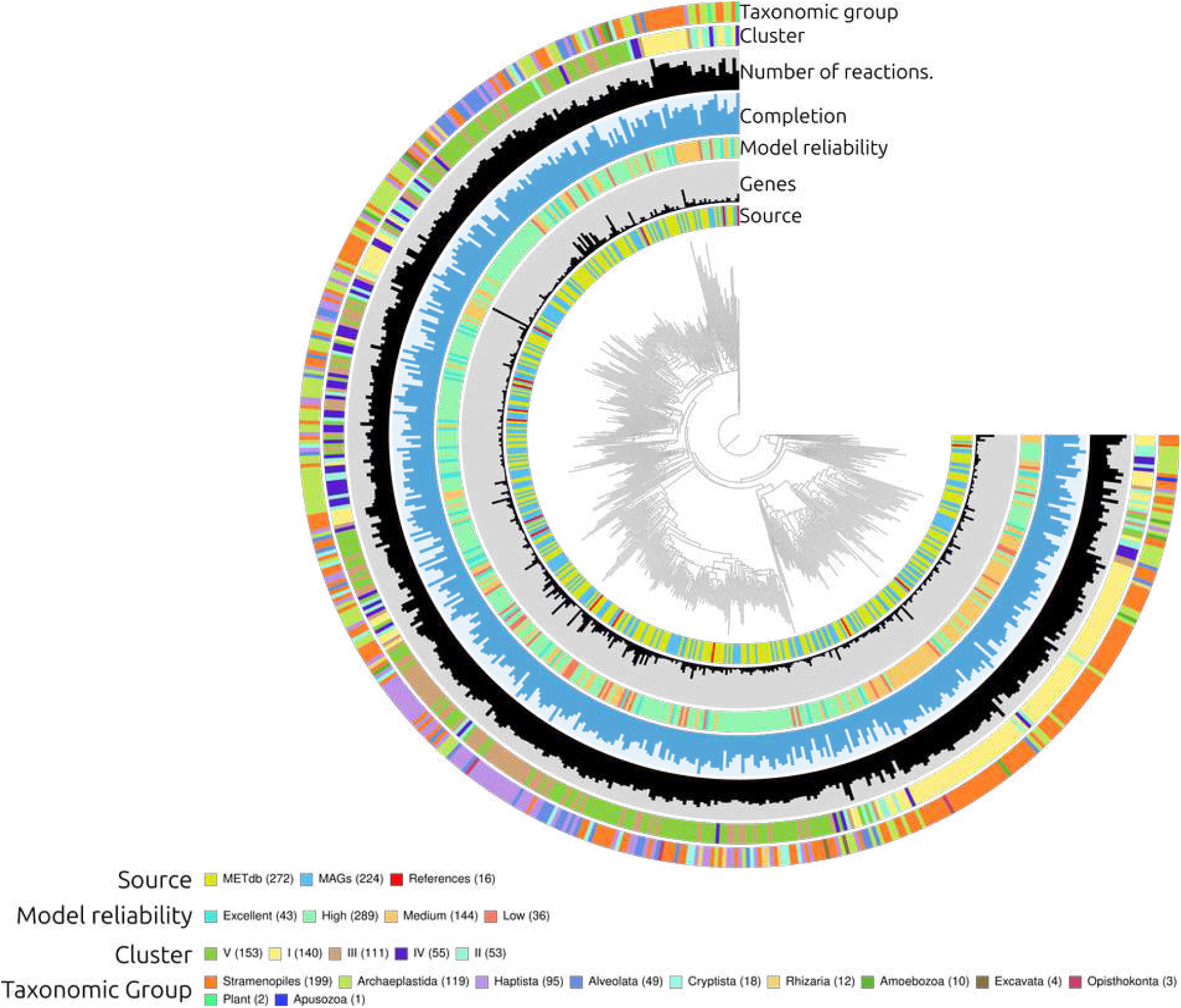
Main characteristics of PhotoEukStein derived GSMs. Central dendrogram represent Jaccard distance between GSMs reactions composition. Inner circle indicate the source of the sequence supporting the GSMs. Model reliability is defined as: Excellent: ≥ 75% completion and < 10% gapfilled reactions (reactions imported without direct genetic evidence) ; High: ≥ 50% completion and < 20% carved reactions ; Medium: ≥ 25% completion and < 30% carved reactions ; Low: the rest. See Extended data for further details. Completion is the Busco-based evaluation of genome completion (Manni 2021) (see Supplementary Table S2 and Online Methods). Number of reactions in the reconstructed GSMs (see Supplementary Table S2 and Online Methods). Clusters as defined in Supplementary Figure S7. Taxonomic groups are from (Delmont 2022) and reported in Supplementary Table S2. For the sake of readability, 10 Amoebozoa, 4 Excavata, 3 Opisthokonta and 2 Plants are displayed as “Others” in the Taxonomic supergroups ring.

**Figure 3:**
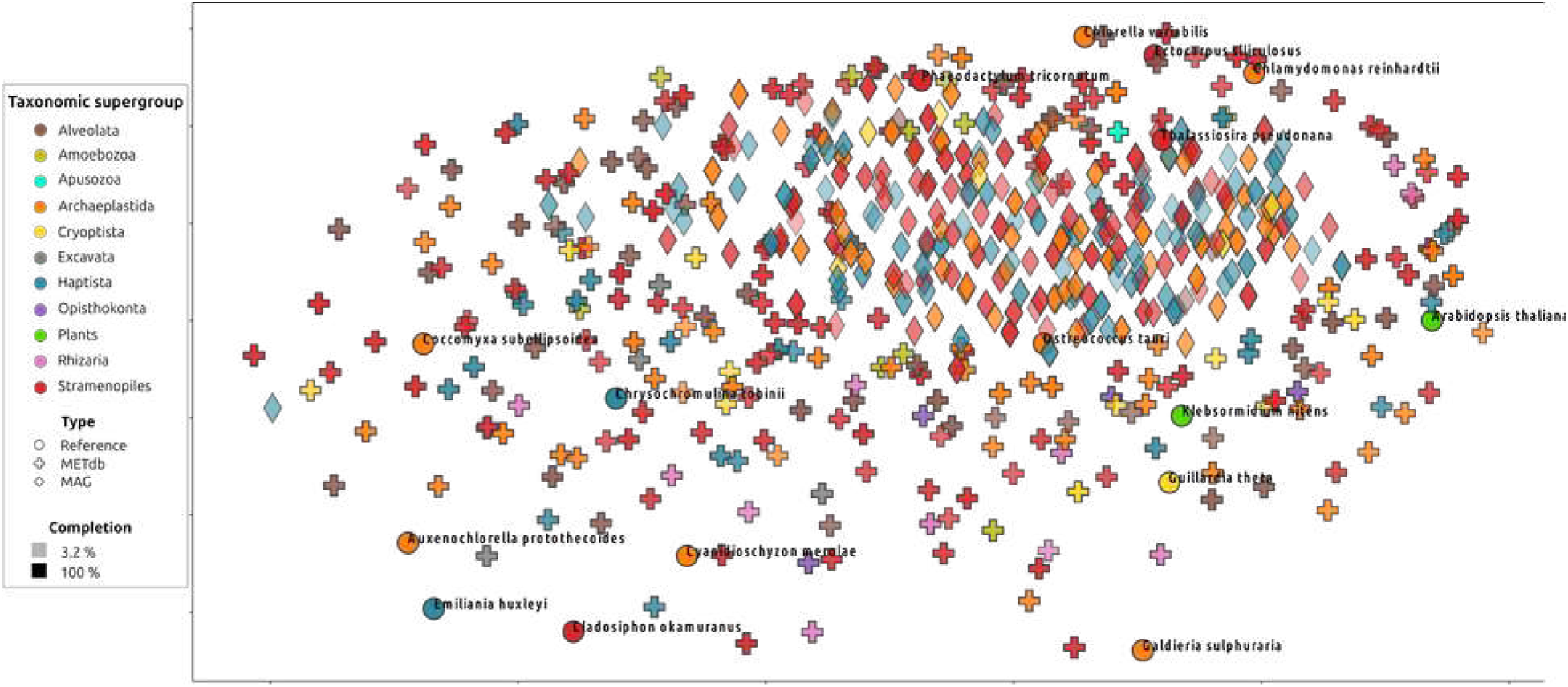
UMAP representation of diffusion map analysis of the 549 GSMs topology (see Methods). MAGs are symbolised by diamonds, METdb by crosses, and reference genomes by circles. Colours indicate the taxonomic groups of each supporting genome, while transparency represents Busco-based genome completion estimation.

We showed a global spread of PhotoEukStein GSMs in the metabolic topological space, indicating that these models are globally functionally diverse but also specific to each genome or transcriptome they are built upon. Moreover, despite the wide breadth of taxonomic diversity covered by these organisms, there needs to be evidence of the structuration of the metabolic connectivity space based on the taxonomy. As diffusion map analysis captures the connectivity between reactions, this observation suggests that the taxonomy does not critically influence the metabolic circuitry of organisms. This result indicates how the various organisms mobilise their metabolic capabilities to produce biomass. These two visions of metabolic behaviour are similar to the genotype and phenotype (i.e., the difference between the potential and its realisation in a given set of conditions).

### Estimation of metabolic traits of 549 genome-scale metabolic models

To further evaluate the ability of PhotoEukStein-based GSMs to respond to environmental changes and, therefore, to capture an organism’ responses to environmental variability, we applied combinations of metabolites fluxes complementing the reference growth medium (Supplementary Table S5) and evaluate GSMs-predicted growth rate variations (Figure 4). In all cases, the predicted growth rate increased compared to the reference medium (Figure 4, Supplementary Table S6). As the metabolites in the permuted pool can fuel a wide range of reactions, their addition increases the usability of possible metabolic routes to produce biomass. However, the various models do not exhibit the same response profile depending on the combination of added metabolites, and we can define groups of models sharing similar profiles, hence defining functional traits (Supplementary Figure S7). Interestingly, growth profiles correlate poorly with taxonomy or genome-wide gene content, similar to what have already been described for Bacteria ^46^.

**Figure 4:**
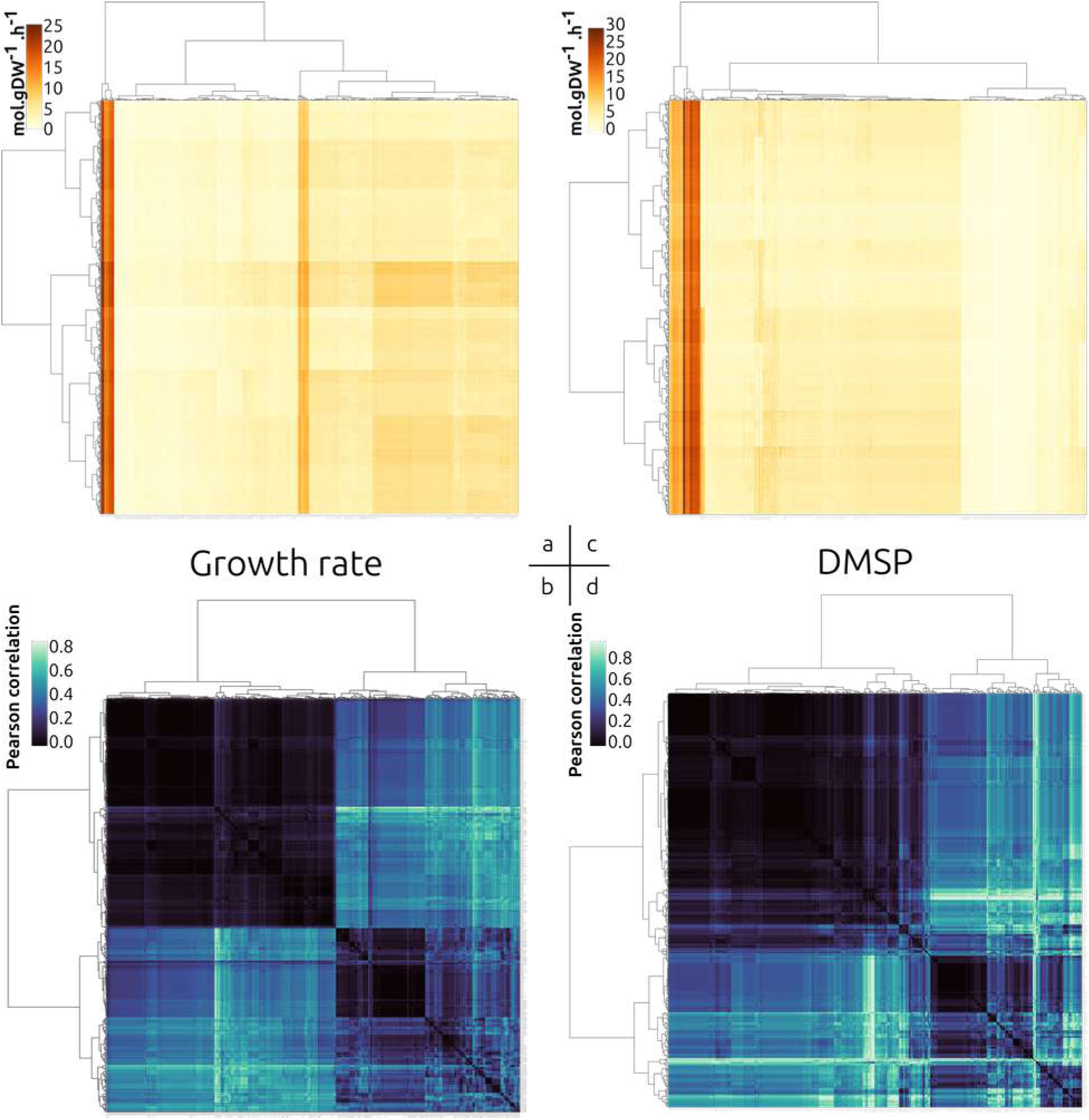
Metabolic niche exploration. The 549 GSM models are exposed to medium modification by systematic permutations of 1 up to 9 extra metabolites (listed in Supplementary Table S4 and S5), and growth rate is computed for each condition and represented in panel a). The GSMs (in columns) are clustered based on their growth rate variation across the 1023 permutations (in lines). b) represents the clustered correlation matrix of growth rates modification profiles across the 1023 permutations. c) DMSP production rate as computed for the 337 GSMs producing that molecule for the same metabolic niche permutations as panel a, and d) shows the clustered correlation matrix of DMSP production variation under the 1023 permutations.

Beyond of the sole growth rate, GSMs can define the functional traits of a given metabolite. Among the list of metabolites of interest for the global carbon cycle ^47^, we considered the 387 GSMs able to produce dimethylsulfoniopropionate (DMSP). This molecule plays many important roles in marine life, including osmolyte, antioxidant, predator deterrent, and cryoprotectant for phytoplankton, as well as a reduced carbon and sulphur source for marine bacteria ^48^. It also produces the climatically active gas dimethyl sulphide (DMS), the primary natural source of sulphur in the atmosphere ^49^. We computed the amount of DMSP produced in the various conditions tested (Figure 4c and Supplementary Table S7). Based on predicted DMSP production values in standard medium (iteration 0 in Supplementary Table S7), two classes for the DMSP producers discriminate, with a first group with DMSP production rates < 3 mol.gDW^-1^.h^-1^ and a second group with DMSP production rates > 6 mol.gDW^-1^.h^-1^ (reported in Supplementary Table S2 as low and high).

This high-rate producers group appears on the left side of Figure 4c. There is no evidence of taxonomic-based enrichment in either groups (high producers contains 8 Alveolata, 4 Archaeplastida, 1 Cryptista, 1 Excavata, 2 Haptista, 6 Rhizaria, 7 Stramenopiles, while low producers group encompasses 33 Alveolata, 3 Amoebozoa, 1 Apusozoa, 56 Archaeplastida, 15 Cryptista, 73 Haptista, 3 Opisthokonta, 5 Rhizaria and 164 Stramenopiles). This observation aligns with what is already reported for DMSP production in eukaryote phytoplankton ^50^. Similarly as for the growth rate variations, we observe that different GSMs respond to medium changes with various patterns. When considering these response profiles, we can define two very differently responsive groups of GSMs that, once again, do not follow taxonomy discrimination. This distinction is the result of underlying functional traits essential for ecosystem model.

N-limitation can lead to increased DMSP production in many DMSP-producing algae and plants, resulting in higher sulphur incorporation relative to nitrogen incorporation ^51–53^. It has also been proposed that DMSP production can act as an overflow mechanism for excess reduced-carbon and sulphur compounds ^51^. The overflow mechanism can be seen as a response of the cell under conditions of unbalanced growth, producing and discarding compounds to ensure the continuation of other metabolic pathways, allowing continued sulphate assimilation even under nitrogen-limited conditions. We therefore explore the behaviour of PhotoEukStein-derived GSMs under nitrogen depletion, with a focus on DMSP fluxes (Figure 5a). For this numeric experiment, NO_3_ ^-^ is the only available nitrogen source in the environment (Figure 5a).

**Figure 5:**
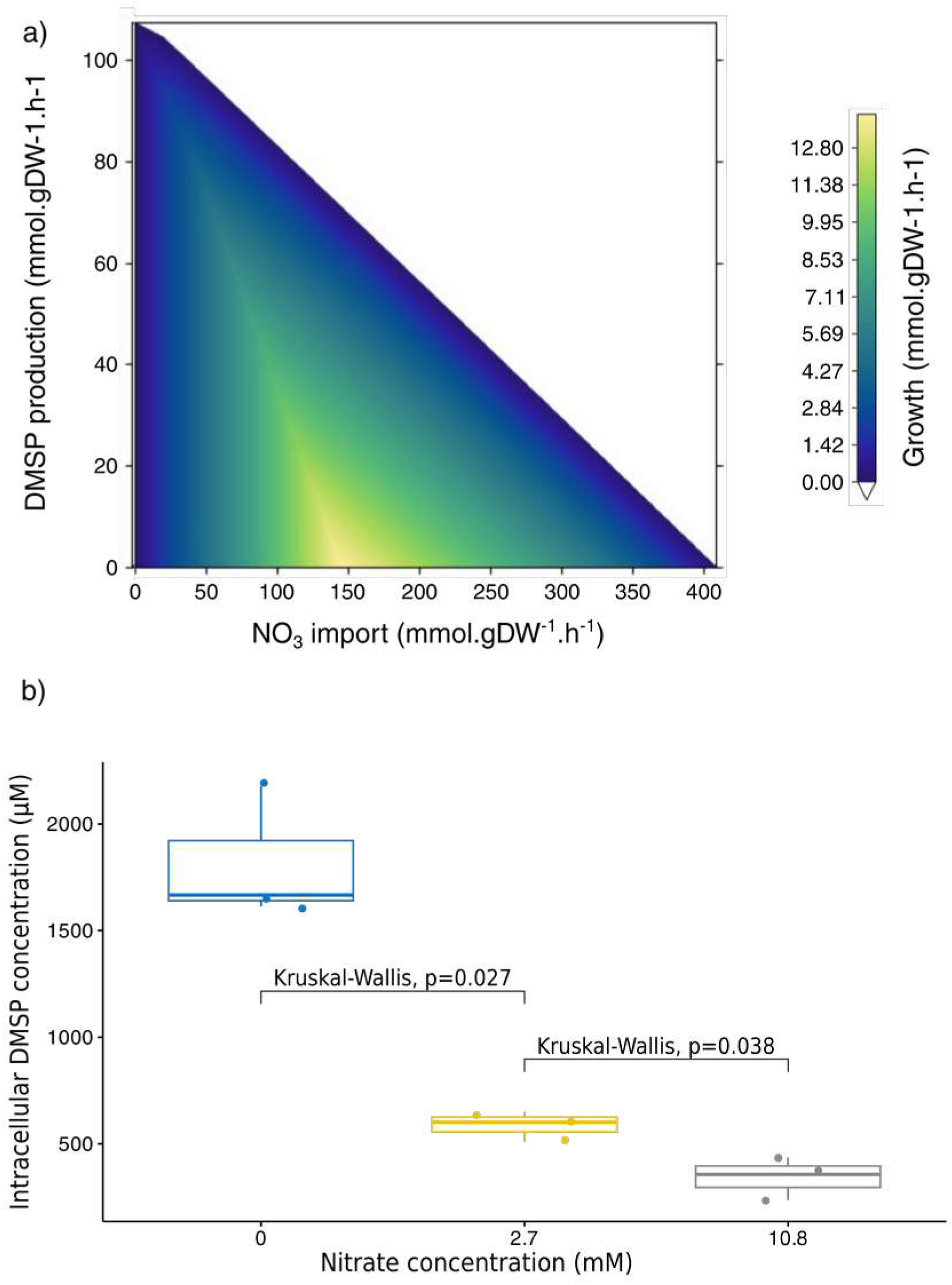
*P. tricornutum* intracellular DMSP concentration under nitrogen deprivation stress a) as predicted from PhotoEukStein-derived GSM model and b) as observed in-vivo after 40 hours of cultivation in normal, nitrate-reduced or nitrate-depleted conditions (see Supplementary Table S10 and the Material and Methods section 6 for details). In a) the solutions space was projected on the following three axes: NO_3_^-^ uptake flux, DMSP secretion, and growth rate.

A quasi-linear relationship is observed between the reduction of the NO_3_ ^-^ uptake flux and the DMSP production. The biomass flux is maximal (about 13 mmol.gDW^-1^.h^-1^) when the system uptakes NO_3_ ^-^ with a rate of 150 mmol.gDW^-1^.h^-1^. When lowering nitrate uptake down to 0, the model predicts that in order to maintain growth, the production of DMSP has to increase. The less nitrogen is imported, the less growth can be maintained and the more DMSP is exported, in agreement with the shunt hypothesis to conserve nitrogen - carbon and sulphur stoichiometry in the system.

We conducted an in vivo experiment to verify this predicted behaviour. Three *Phaeodactylum tricornutum* separate cultures in exponential growth conditions were exposed during 40 hours to decreasing concentrations of nitrate as sole source of nitrogen in the medium (10.8 mM – normal nitrate concentration, 2.7 mM and 0) and intracellular DMSP concentration was then measured (Figure 5b, Supplementary Table S10. See Materials and Methods section 6 for details). As predicted by the model, and conform to previous observations ^51,53^, we observe a statistically significant increase in DMSP intracellular concentration with the decrease of nitrate availability in the medium. A residual growth is maintained even at 0 mM of nitrate, probably due to the delay in residual nitrogen consumption and intracellular recycling.

Overall, we observed the relationship between nitrogen availability in the culture medium and intracellular DMSP concentration, as modelled by PhotoEukStein-based metabolic model for *Phaeodactylum tricornutum*. Not only unicellular phototrophs eukaryotes GSMs can describe global phenotypic properties, but can also, to a certain extend, predict precise biologic features, and therefore be a tool to determine functional features of organisms.

## Discussion

While bacterial and archaeal metabolisms have been extensively studied, more efforts have yet to be devoted to eukaryotes, and even more for marine multicellular species. With the growing number of available environmental and isolate genome data, the repertoire of available MAGs and transcriptomes representing planktonic eukaryote species distantly related to reference organisms is franticly expanding. However, while they cover a broader diversity than the well-studied references, there needs to be more efficient ways to study their biology ^54^.

We propose PhotoEukstein as the first meta-metabolic model of unicellular phototroph eukaryotes for a fast and efficient top-down derivation of GSMs, applicable to genomes and transcriptomes. Its development and curation involved integrating diverse experimental data and literature sources and extensive manual curation and refinement, resulting in a model that captures a broad range of metabolic processes and interactions. We efficiently applied it to a collection of taxonomically diverse environmental genomes and transcriptomes covering a wide range of yet barely functionally described marine eukaryote unicellular planktons. We have shown that growth rates predicted from these GSMs are highly comparable for the three reference organisms and with the ones obtained from in-vitro measurements. While automatically derived GSMs may not suffice to cover all fine metabolic behaviours of species (such as complete secundary metabolism), we shown that PhotoEukStein-derived models support easily the comparison with more dedicated tools, and provide a solid asis for species specific improvement by experts, thus enabling a wider community to explore biological features of unicellular phototroph eukaryotes. Therefore, PhotoEukStein represents an essential step towards understanding and modelling phototrophic eukaryotes’ metabolism, physiology, biogeochemistry and ecology. We propose a valuable resource of 549 new metabolic models for researchers, paving the way for an in-depth ecosystemic exploration of plankton communities from viruses to single-cell phototrophs. Moreover, PhotoEukStein can easily be extended to incorporate new metabolic knowledge to cope with the development of eukaryote phototrophs unicellular organisms studies, either through identifying or accumulating reference protein sequences associated with a given reaction or the description of new metabolic reactions and related protein sequences. Deriving PhotoEukStein-based GSMs from new genomes or transcriptomes requires minimal computational resources or time-consuming expertise, allowing it to cope with the rapidly growing repertoire of environmental genomes.

Metabolic models are well suited to represent the metabolic phenotype of microorganisms ^22^. They may provide a better specific delineation of functional traits distribution across species than solely considering taxonomy or the presence/absence of particular genes. On one hand we have annotation based distance that follows phylogenetic distance ^8^, on the other hand we have laboratory experiments that assess phenotypes without finding a correlation with phylogenetic distances ^46^.

Similarly to the later, our results show that a robust phylogenetic signal affects the composition of the metabolic reactions profile (Figure 2 and Supplementary Figure S3). In contrast, no such signal is detected in functional/phenotypic GSMs clustering (Figure 3). It results that closely related organisms with similar repertoire of metabolic reactions may display dissimilar functional profiles, and (inversely) that distantly related organisms might exhibit metabolic similarities. It relates to the consideration that metabolic reactions act together to form biological functions. The profiling of organisms for each given functional trait will generate specific classifications that cannot a priori be reduced prior to taxonomy or the presence/absence of a gene. Understanding the biological functions of organisms involves deciphering their metabolic capabilities, and using GSMs for this purpose could be the most effective even when only environmental genomic data are available.

Being able to derive functional GSMs for unicellular phototrophs eukaryotes, even from environmental omics data, provides a unique way to assess their biological phenotype *per se* as it differs from the sole identification of functional genes. These features are new observations, or semantic traits that arise from genomics descriptions. For example, scrutinising the variability of metabolic fluxes and metabolite exchanges through the study of metabolic niches may allow differentiating allocation of cellular resources to resource acquisition, defence, signalling, and other survival needs ^55^, as well as community metabolic interactions as considered in the phycosphere ^56^ or the holobiont ^57^ concepts. Therefore, the ability to systematically derive GSMs for unicellular eukaryote phototrophs, as is already the case for heterotroph prokaryotes, is an essential step toward a global description of phenotypic biodiversity and ecosystems modelling. In particular, we advocate for considering PhotoEukStein and derived GSMs as a resource for emphasising better omics-driven phenotypes estimations and functional trait definition that might benefit to future ocean system models ^58,59^.

## Acknowledgements & Funding

Our survey was made possible by the sampling and sequencing efforts of the *Tara* Oceans Project. We are indebted to all who contributed to these efforts and other open-source bioinformatics tools for their commitment to transparency and openness. *Tara* Oceans (which includes the *Tara* Oceans and *Tara* Oceans Polar Circle expeditions) would not exist without the leadership of the *Tara* Oceans Foundation and the continuous support of 23 institutes (https://oceans.taraexpeditions.org/). We also acknowledge the commitment of the CNRS and Genoscope/CEA. Some computations were performed thanks to the TGCC computing facility in France. This study was supported in part by FRANCE GENOMIQUE (ANR-10-INBS-09). M.B. was funded by the Doctoral School “Structure and Dynamics of Living Systems” of Paris-Saclay University. This research was co-funded by the European Union (GA#101059915 – BIOcean5D). Views and opinions expressed are however those of the author(s) only and do not necessarily reflect those of the European Union. Neither the European Union nor the granting authority can be held responsible for them. We also thanks Jeanne Got for technical discussions. This article is contribution number XXX of Tara Oceans.

## Licence

© The Authors. This manuscript (including its supplementary informations) is under the Creative Commons CC-BY-NC license (http://creativecommons.org/licenses/by-nc/4.0/)

## Data availability

PhotoEukStein is available at https://www.genoscope.cns.fr/PhotoEukStein/ Supplementary tables are available at https://www.genoscope.cns.fr/PhotoEukStein/ PhotoEukStein installer and collection of generated models are available at Zenodo under doi 10.5281/zenodo.13862627

## Authors contributions

M.B., A.R., D.E. and E.P. designed the research, M.B. build, curated and applied PhotoEukStein, M.B. and A.R. validated PhotoEukStein, designed numerical experiments with inputs from D.E, and generated the data, C.O. and A.T. cultivated *P. calceolata*, *P. tricornutum* and *T. amphioxeia*, S.B. analysed experimental and predicted growth rates data, N.P. and A.P. performed metabolites extraction and LC/MS/MS analysis, all authors analysed the results, M.B. and E.P. wrote the manuscript with significant inputs from all authors.

## Material and Methods

### 1. Constraint-based metabolic modelling at genome-scale

Metabolic networks contain the metabolic capabilities encoded in the organism’s genome. Indeed, from the genome of a specific organism, it is possible to predict the encoded genes and thus identify the corresponding enzymes and their associated metabolic reactions. Metabolic reactions are the set of life-sustaining chemical transformations in organisms. They allow organisms to grow, reproduce, maintain their structures and respond to their environments. Because the products of some reactions are the substrates of others, the reactions are interconnected by what are called metabolites. Metabolic networks are modelled in order to study the physiology of the relevant microorganism. In particular, metabolic models are used to infer reaction rates, also known as fluxes, without using kinetic parameters. A metabolic model is formally described by its stoichiometric matrix *^S^*, where the rows correspond to the metabolites, the columns correspond to the reactions considered in the metabolic network. The entries are stoichiometric coefficients which are negative if the metabolite is a substrate, positive if the metabolite is a product and null if the metabolite is not implicated in the reaction. Assume there are *^m^* metabolites *^M^_i_* to *^M^_m_* and *^n^* reactions *^r^*_1_ to *^r^_n_*. We note respectively *^v^*_1_ to *^v^_n_* the fluxes of *^r^*_1_ to *^r^_n_*. Let consider the stoichiometric coefficients of *^M^_i_* in reactions *^r^*_1_ to *^r^_n_*. The change over time of the *^M^_i_* concentration is given by the mass-balance equation:

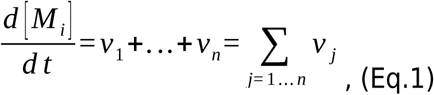

Using a vector notation, the above equation can be written as:

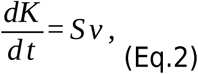

where *^v^* stands for the fluxes vector, and *^K^* is the metabolites concentration vector. A metabolic network is formally described by its stoichiometric matrix *^S∈Rn,m^* describing the relationship between the *^n^* metabolites and the *^m^* reactions. The entry is the stoichiometric coefficient of the metabolite *^M^_i_* in the reaction *^R^ _j_*. By convention it is negative if the metabolite is a substrate, positive if the metabolite is a product and null if the metabolite is not implicated in the reaction.

In general, the rate of reactions depends on metabolites concentrations and kinetic parameters, such as temperature or pH. Determining these parameters and the function of reaction rate are complex experimental tasks. Moreover, these parameters are generally very sensitive to biochemical conditions, so *in vitro* determinations may not correspond with *in vivo* values ^1^.

Thus solving Eq.2 is a daunting task for genome-scale systems. When analysing metabolic networks using constraint-based approaches, we assume that organisms are homeostatic, keeping internal concentration as constant as possible using regulation ^2^. Thus, the rate of formation of internal metabolites is equal to the rate of their consumption. The system is then considered in a quasi-stationary state ^2^, leading to:

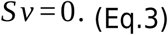

In addition to this system of linear constraints, we also consider thermodynamic constraints on fluxes. Fluxes can be positive or negative. For inner reaction, a positive flux means the reaction is occurring in its forward direction, whereas a negative flux means it is happening in reverse. All fluxes must satisfy an inequality like:

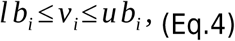

where *^l b^_i_* represents the lower bound of the flux, and *^ub^_i_* represents the upper bound of the flux. Fluxes are expressed in mole of product formed by gram of dry weight of the considered organism by the hour (mol.gDW- 1.h- 1). Knowledge on the reaction direction and reversibility can also be encoded in those inequalities. For instance, if the reaction is known to be direct and irreversible, the flux cannot be negative. Eq. 4 becomes:

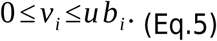

These equations result in a model described as a set of constraints. Altogether, Eq.3 and Eq.4 form a constraint-based metabolic model (CBM) of the corresponding organism. A CBM at a genome-scale is called a Genome-Scale Metabolic Model (GSM). It can be resumed in the system:

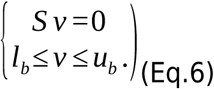

All solutions of *^v^* satisfy all constraints : 1) the steady state equation, and 2) the thermodynamic constraints, thus defining a steady- state flux space. This “flux space” may be further analysed through several state-of-the-art approaches., the reader may wish The reader may refer to ^3–5^ for a detailed review of these methods. A metabolic network and its associated GSM allows us to explore the metabolic phenotype of an organism ^2^.

Exchange reactions facilitate the continuous supply of metabolites from and to the media. They are responsible for the uptake or secretion of metabolites. For boundary reactions, a positive flux means metabolite secretion into the environment, whereas a negative flux means metabolite uptake.

Finally, modellers developed fictive reactions to model the growth rate of organisms ^6^, and among them is the biomass reaction. This reaction encompasses the needs of the modelled system (nucleotides for DNA, RNA, amino acids for proteins, lipids, carbohydrates…), but also the energy cost of cellular division or cell maintenance.

## 2. PhotoEukStein : generic model reconstruction

### Metabolic network of PhotoEukstein

PhotoEukStein was built by merging available biochemical and genomic information of 16 autotrophic eukaryotes (Figure 1.A). In biological databases and data integration, using different identifiers for the same entity can create confusion and make merging data from various sources difficult. Thus, identifiers of reactions and metabolites are homogenised to the same namespace using MetaNetX ^7^ and manual curation. Duplicated entities are then removed. To enable seamless integration of the metabolic pathways of 16 different organisms into a single supraorganism, all enzymatic reactions were assumed to occur in a single compartment (with an exception, see below). All network reactions are mass-balanced to predict reactions fluxes without relying on kinetics data (see Eq.6 above). Chemical formulae have been added to all metabolites using MetaNetX, manual curation and prediction algorithms. Duplicated reactions (*i.e.* those that propose similar metabolic transformations but have different identifiers) are deleted. Some are modified based on the literature. When a reaction was not balanced, had no associated metadata, no associated genes, or genes found only in *Arabidopsis or Okamuranus*, the reaction is mostly deleted.

### Constraint-based metabolic model of PhotoEukStein

According to the general protocol in ^8^, the ’draft’ model was then curated manually to generate a functional CBM of eukaryotic-algae metabolism.

To maintain the stationary state of the network, 674 sink reactions (SK) were added for metabolites consumed but never produced (with hard-constraint on the uptake flux : ^–0.5≤v^SK ≤0), and 1033 demand reactions(DM) for those produced but never consumed ( 0 *≤ v _DM_ ≤* 10 0 0). Sink and demand reactions are particular reactions that allow us to keep active metabolic pathways of which some knowledge is missing (they allow an active flux in 2,554 reactions. Indeed, if all SK and DM were blocked, they would turn off 2,554 network reactions. As new enzymatic reactions are discovered, the number of sinks and demand reactions may be reduced in future versions of PhotoEukStein

The directionality of some reactions has been corrected. For example, heuristic rules have been applied to avoid futile cycles or proton gradients that could generate out of nowhere reactions that consumes ATP (except the respiration pathway), ABC transporters and proton pumps are irreversible. Blocked reactions and orphan metabolites were deleted to avoid false-negative analysis regarding gene deletion on flux redistribution. The photosynthetic system of PhotoEukStein is based on iLB1034 ^9^. Adding a pseudo-thylakoid and a chloroplast allows for a spatial organisation that couples the photosynthetic apparatus, chloroplast ATPS, and carbon dioxide fixation by RuBisCo with light absorption, and powers the growth rate (see Extended Informations).

PhotoEukStein encompass 5,831 metabolites and 11,229 reactions. Two types of reactions are distinguished : 2,067 boundary reactions (including 360 exchanges reactions, 674 sink reactions, 1,033 demand reactions), and 9162 internal biochemical transformations. The meta-model has 15 biomass objective functions : one autotrophic BOF from *C. variabilis* ^10^ ; 3 (one autotrophic, one hetereotrophic and one mixotrophic) from *C. reinhardtii* (iRC1080, BiGG, ^11^) ; 11 (two for biomass production during light or dark, and many for specific class of metabolites as DNA, RNA, lipids, carbohydrates production) from *P. tricornutum* (iLB1034 ^9^). The BOF of iLB1034 is used for this manuscript.

### Gene-Protein-Reaction rules

For each internal reaction in the curated universal model, we identify all those equivalent in the input models (i.e., duplicates) to recover the maximum number of logical gene conjunctions (monomeric, oligomeric, isoenzymes or multifunctional enzymes). Thus, 7,599 PhotoEukStein reactions (out of 9162) are associated with 20,468 protein sequences from reference genomes by their respective logical associations. Protein sequences are mostly retrieved from NCBI, UniProt, Diatomics and TAIR (*Arabidopsis thaliana* database).

17% of PhotoEukStein’s reactions do not have associated genes either because the reaction is spontaneous or no genes have been found yet to catalyse the reactions. PhotoEukStein can easily be extended to incorporate new metabolic knowledge to cope with the development of eukaryote phototrophs unicellular organisms studies, either through identifying new metabolic reactions, or accumulating reference protein sequences associated with a given reaction. For example, DMSP synthesis from methionine has been shown to occur via transamination pathway in some eukaryotic algae ^12.^ Although some of the models that make up PhotoEukStein (Figure 1.A) had the DMSP synthesis pathway (e.g. *Thalassiosira* ^13^, *Cladosiphon* o*kamuranus* ^14^, or *Phaeodactylum* ^9^, none had a gene associated with the key enzyme of this pathway. However, two genes encoding for this enzyme in eukaryotic algae have been identified: (i) DSYB, and (ii) TpMT2 whose function was confirmed in *T. pseudonana* ^14^. We added 135 sequences to DSYB, and 6 for TpMT2 (from ^12,14,15^) in the protein sequences database of PhotoEukStein. 337 models of the GSMs database can produce DMSP (Supplementary Table S7).

The genomic and biogeochemical information of *Thalassiosira pseudonana* included in PhotoEukStein comes from the PGDB of BioCyc (2012). The constraint-based model used to make the comparisons comes from a recent publication ^16^. Thus, unlike the other GSMs, the reference used to validate the PhotoEukStein-derive model of *Thalassiosira* is not included in PhotoEukStein (therefore its BOF is also different). This may explain the more significant differences in growth rates between the two models in Figure S1. However, the correlation being very high, we can see that the two models adapt to their environment similarly.

## 3. PhotoEukStein validation

### Phototrophic phenotypes of PhotoEukStein

We ensure that PhotoEukStein can grow under photoautotrophic conditions with adapted physiological strategies like the ability to fix inorganic carbon. We expected a coupling between light uptake and CO_2_ uptake from the environment and the underlying synchronisation of photosystem reactions: ATP production by the chloroplastic ATP synthase and inorganic carbon assimilation by the ribulose-1,5-biphosphate carboxylase (RuBisCo).

Under photoautotrophic conditions (cf. Medium section below), we computed a projection of the allowable flux space on critical reactions fluxes ^17^ of PhotoEukStein to scrutinise the fluxes variability and coupling of these essential reactions, and thus assess some photoautrophic phenotypes for PhotoEukStein (see Extended Data).

### Growth rates comparison

To validate PhotoEukStein, we compare our automatically reconstructed models with reference model of the same organism. The organisms were reconstructed with a medium allowing the reference model to grow (see Medium section below).

When computing the niche space of models ^17^, we compare the flux through the biomass reaction for around 10^4^ randomly generated environmental conditions. The environmental condition are composed of fixed fluxes of exchange reactions concerning the following metabolites: CO_2_, photon, SO_4_, NH_4_, NO_3_ or Phosphate. No other constraints were applied to the exchange reactions of the reference models. For the PhotoEukStein derived models, if the exchange reaction exist in the corresponding reference model, the bounds are the same as the reference, otherwise the lower bound is set to 0 except for the exchange reactions concerning H_2_O, H, Mg^2+^, Fe^2+^and Fe^3+^.

### A sampling of the solution space

#### Reaction fluxes correlations

To compare further the models, we applied a sampling procedure of all the allowable solution space of 2 models of *Phaeodactylum tricornutum*. *Phaeodactylum tricornutum* original GSM (iLB31034, ^9^) is composed of 2162 reactions, while PhotoEukStein-derived model (phaeo- photoeuk) is composed of 5366 reactions (Supplementary Table S2). The fluxes constraints of the exchange reactions of iLB1034 have been applied on phaeo-photoeuk. For each model, a sampling procedure have been applied (see https://cobrapy.readthedocs.io/en/latest/sampling.html, OptGPSampler, thinning=10,000, sample=10,000). Blocked reactions are removed, and the shared reactions are considered (1171 reactions). From those, fluxes correlations (Pearson) for each pair of reactions were computed.

Only the reactions with at least 1% of their absolute correlations being higher than 0.2 (and which are shared by the two models), have been kept for the analysis (434 reactions, Figure S2). Python package Seaborn.heatmap has been used for the plot.

#### HexBins

From the 94,178 correlations (434x434/2): upper or lower triangle of correlation matrix), we eliminated (i) the absolute correlations that were lower than 0.025 in both models, and (ii) the 434 correlations of the diagonal, resulting in 69,442 remaining correlations. We compare the values of these correlations with the HexBins. The x-axis corresponds to the correlation values of phaeo_photoeuk, and the y-axis to iLB1034. For each correlation, we plot the result for each model. Python package Seaborn.jointplot has been used for the plot.

## 4. Experimental Validation of PhotoEukStein

We performed growth experiments on a stramenopile (*Pelagomonas calceolata* RCC100) and a cryptophyte (*Teleaulax amphioxeia* AND-0710). Both were grown at 20°C in L1 medium prepared with artificial seawater (ASW), as described in ^18^. Nitrate-free L1 medium was supplemented with NaNO₃ to obtain five nitrate concentrations (3.48, 1.72, 0.882, 0.441 and 0.110 mM). Cultures were grown in flasks under four white-light intensities: 2, 5, 10 and 20 µmol photons m⁻² s⁻¹ for *P. calceolata* and 10, 20, 50 and 100 µmol photons m⁻² s⁻¹ for *T. amphioxeia*. Cells abundance was monitored by flow cytometry (CytoFLEX, Beckman Coulter) using the B690 channel for chlorophyll fluorescence (Supplementary Tables S11 and S12, respectively).

Logistic growth curves were fitted to raw cell density time series for each combination of light intensity and nitrate concentration tested. To focus on the exponential growth phase and avoid bias introduced by nutrient depletion or cell death, only data points up to the observed maximum cell density were retained for each condition. The logistic function:

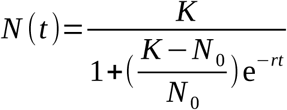

was fitted to the truncated time series, where N(t) is cell density at time t, K is the carrying capacity, N_0_ is the initial cell density, and r is the logistic growth rate (d^−1^) (Supplementary Figures 7 and 8).

## 5. Exploration of PhotoEukStein-derived metabolic models DB

### PhotoEukStein-derived models

#### Tara Oceans eukaryote MAGs resource

MAGs sequences (predicted CDS and their functional annotations) corresponding to Delmont *et al.* 2022 ^19^ were downloaded from the original source https://www.genoscope.cns.fr/tara/

#### The METdb database for eukaryote transcriptomes

METdb is a curated database of transcriptomes from marine eukaryotic isolates that cover the MMETSP collection (new assemblies were performed, combining time points from the same culture in co-assemblies when available) as well as cultures from TARA Oceans ^20^. The database is publicly available and can be accessed at http://metdb.sb-roscoff.fr/metdb/.

#### Identification of phototrophs MAGs or METdb

The subset of phototrophs MAGs and METdb was defined as those encoding proteins with the Chlorophyll A-B binding protein domain (InterPro entry IPR022796)

#### Deriving GSMs from PhotoEukStein

CarveMe can be easily installed using the pip package manager. Additionally, the Diamond package and IBM CPLEX Optimizer must be installed manually (see https://carveme.readthedocs.io/en/latest/installation.html) .

To use PhotoEukStein with CarveMe, one need to download

- (1) the generic model,
- (2) the Gene-Protein-Reactions associations files,
- (3) the protein sequences database and
- (4) the media file. The sbml.py from the Cobra package need to be changed to support the reading of identifiers from BioCyc. Please read the associated
- (5) README file for more information on how to proceed.

The organisms were reconstructed with a medium allowing the reference model to grow (see supp Mat medium). For the reconstruction of PhotoEukStein-derived models the following command have been used: carve -d -v path/to/input/fasta.faa --universe photoeukstein --gapfill medium_name --output path/to/output/model.xml

« medium_name » is set to « phaeo » for the whole model reconstruction except for the computation of growth rate comparison with references for *Chlorella variabilis* (medium « chlorella »)*, and Thalassiosira pseudonana* (medium « thalassio »)

#### Compositional analyses

The presence/absence of 9648 reactions (those mobilised by model resources) were used to hierarchically cluster (Euclidean distance, Ward distance) both GSMs (lines) and reactions (columns). Genome source, Taxonomic group, model quality score (as defined in Extended Data), and metabolic cluster (as described in Figure S5) are shown for each GSM. The frequency of reaction appearance is listed in Figure S4. We used the UMAP algorithm ^21^ to visualise how the presence/absence of the 9648 reactions (those mobilised by the model’s resources) helps structure the models together. A k-means clustering was used to identify the clusters used in Supplementary Figures S4 and S5.

#### Topological analysis

We analysed the dataset through the algorithm developed in ^22^ . We used the GSM reconstructed with their sink reaction ; however, only the internal reactions were considered. All the diffusion variables were then normalised, and we used the UMAP algorithm for better visualisation ^21^.

#### Functional analysis

To assess an organism’s functional phenotypes (towards new functional traits), we consider a primary medium (Supplementary Table S4) and add a set of new nutrients (up to 9, Supplementary Tables S4 and S5). For each condition, we maximise the flux through the biomass reaction. The set of new nutrients results from using the itertools combinations algorithm in the Python library on the list of considered nutrients (Supplementary Table S5).

#### Anvio representations

All Anvio-based representations (Figures 1 and 2) were generated using Anvio version 7.1 (http://anvio.org/) ^23^ from data available in Supplementary Tables S2 and S3.

#### Growth medium for GSMs

We have retrieved the respective mediums of each reference model (*Chlorella*, *Thalassiosira*, *Phaeodactylum*). To achieve this, we have performed an FVA analysis. Then, if the flux interval indicates values less than 0 for each exchange reaction, the metabolite can be imported into the system. It is, therefore, part of the medium of the organism considered.

#### GSMs evaluation using MEMOTE

PhotoEukStein-derived and reference GSMs quality where evaluated using the MEMOTE tool ^24^ (version 0.17.0) using the Cplex solver from IBM, with the following command line: memote report snapshot --solver cplex.

## 6. DMSP production by Phaeodactylum tricornutum

### Growth conditions and sample collection

*Phaeodactylum tricornutum* (Pt1/RCC2967) was cultured in three replicates in f/2 medium on artificial sea water (ASW - NaCl 0.46M, MgSO_4_ 27 mM, MgCl_2_ 28 mM, CaCl_2_ 10 mM, KNO_3_ 9.9 mM, KH_2_PO_4_ 0.514 mM, NaHCO_3_ 0.476 mM) complemented with: NaNO_3_ 0,88 mM, NaH_2_PO_4_ 36.2 µM, Na_2_Si0_3_ 106 µM, f/2 vitamins and f/2 trace metals ^25^ during 7 days in a growth chamber at 20°C. Culture were maintained in plastic flasks shaken at 150 rpm under a photon irradiance of 80 µmol.m^-^².s^-1^ provided by a cold-white led light with a 12-h dark / 12-h light photoperiod. Cell density reached 12.6x10^6^.ml^-1^, as determined by flow cytometry (CytoFlex, Beckman Coulter).

The culture was then split in three, and after centrifugation (3,000 g, 15 min, room temperature), cells from each split were recovered and resuspended at 6x10^6^.ml^-1^ either in a fresh f/2 medium (10.8 mM NO_3_ ^-^) or in a modified f/2 ASW medium (f/2-N medium) in which KNO_3_ was replaced by 9.9 mM KCl -- (0 mM NO_3_ ^-^) or in a mix of respectively ¼ – ¾ of each (2.7 mM NO_3_ ^-^).

DMSP production was evaluated from a volume of culture containing 10^8^ cells collected 40h after the onset of starvation by filtration onto a 47 mm PTFE filter (JH Omnipore, 0.45 µm) and immediately processed for metabolite extraction.

### Metabolite extraction and LC/MS/MS analysis

Metabolite extraction was adapted from ^26^. Briefly, after filtration, the filter containing the cells was placed face-down in a glass dish containing 5 ml of a cold mixture of H_2_O/methanol/acetonitrile (1/3/1) and submitted to a mild sonication in an ultrasonic bath (Bronson 2510 Ultrasonic Cleaner) for 10 min at 4°C to remove the cells from the filter. The liquid containing the cells was transferred into cryogenic vials and underwent 3 freeze/thaw cycles in liquid nitrogen/65 °C water to fully break the cells and extract the metabolites. The debris were removed by centrifugation (20,000 g, 10 min, room temperature). The supernatants were dried and dissolved in 300 µl water, filtered on 0.22 µm (polytetrafluoroethylene; AcroPrep Advance, Pall) and finally diluted to 1/2000 in a solution composed of 80% acetonitrile and 20% 10 mM ammonium bicarbonate (pH 8.0) before LC/MS/MS analysis.

DMSP (C_5_H_11_O_2_S, m/z 135.04748) was detected by LC/MS/MS in the positive ionization mode using Thermo Scientific™ Dionex™ Ultimate 3000 Rapid Separation (3000RS) liquid chromatographic coupled to ultra-high resolution Orbitrap Elite hybrid mass spectrometer from Thermo Fisher (Courtaboeuf, France) equipped with an heated electrospray ionization (HESI) source.

HPLC separation was achieved on an Atlantis Premier BEH Z-HILIC Column 100 × 4.6 mm^2^; 5 μm (Waters, U.S.A.) thermostated at 40°C. The mobile phase flow rate was set at 0.5 ml.min^-1^ and 4 µl were injected. Mobile phase A consisted of 10 mM ammonium bicarbonate, pH 8.0 and mobile phase B consisted of acetonitrile. The gradient started at 80% B for 1 min followed by linear gradient to 40% B for 7 min, and remained at 40% B for 2.5 min. The system returned to the initial composition (80% of solvent B) in 2 min and re-equilibrated under these conditions for 10 min.

Mass spectrometry analyses were conducted with the following parameters : electrospray voltage 4.20 kV, sheath gas and auxiliary gas flow rates 60 arbitrary units (a.u.) and 44 a.u., respectively, the source heater temperature 250°C and the capillary temperature 275°C. The mass resolution was set to 60,000 m/Δm. Mass spectra were acquired over an m/z range from m/z 50 up to m/z 600.

Raw data were analysed using the Qual-browser module of Xcalibur version 2.2 (Thermo Fisher Scientific, Courtaboeuf, France).

Intracellular DMSP concentration of *P. tricornutum* was computed based on a mean cellular volume of 32,6 µm^3^ ^27^. Resulting values of intracellular DMSP concentration variation following nitrate availability in the medium are indicated in Supplementary Table S10.

Statistical significance of intracellular *P. tricornutum* DMSP concentration variation under nitrogen depletion was evaluated with a one-way ANOVA test.

Fig 5a was obtained through the projection of the metabolic niche on the axes NO_3_, DMSP and Growth, as done in ^17^. No threshold on the growth value was applied, *i.e.* no death rate was used, as we are here exploring the production of DMSP that can occur at a growth of 0.

## Extended Data

### 1 - PhotoEukStein validation

In an epistemological context, model validation and sensitivity analyses are critical steps to ensure the robustness and reliability of the model’s predictions. The validation process generally consists of comparing the model’s predictions with observed data, the literature or other reliable models, and thus assessing the model’s ability to reproduce known phenomena. Then, one can use this model to test new hypothesis and predict future outcomes for which one does not yet have empirical values. When a generic model is converted to ready-to-use organism-specific models using CarveMe, the whole manual curation and relevant structural properties are preserved (Machado 2018).

We ensure that PhotoEukStein can grow under photoautotrophic conditions (see Medium in Material & Methods section) with adapted physiological strategies. Indeed, the ultimate goal of photoautotrophic organisms is to use light energy to convert water and carbon dioxide into oxygen and energy-rich organic molecules such as glucose or starch. Therefore, we expected a coupling between light uptake and CO_2_ uptake from the environment, and the underlying synchronization of photosystem reactions, ATP production by the chloroplastic ATP synthase, as well as inorganic carbon assimilation by the ribulose-1,5-biphosphate carboxylase (RuBisCo being the key enzyme of Calvin cycle). In order to validate the basic internals of PhotoEukStein, and more specifically the phototrophy-associated reactions, we computed a projection of the allowable solution space (Régimbeau et al., 2022) on these key photoautotrophic reaction fluxes to determine their distribution and couplings. The more photons enter the system, the more the photosystems are stimulated with a synchronization of the two photosystems (figure A). We also see that the ATP production by chloroplastic ATPS is coupled to the photosynthetic activity and fuels the growth reaction (figure B). This ATP production is also coupled to CO_2_ uptake and the activity of RuBisCo (figure C). Overall, the uptake of photon into the system stimulates the photosystem apparatus (PSII, PSI) and empowers ATP production. The ATP allows the CO_2_ fixation by RuBisCo and fuels the biomass production.

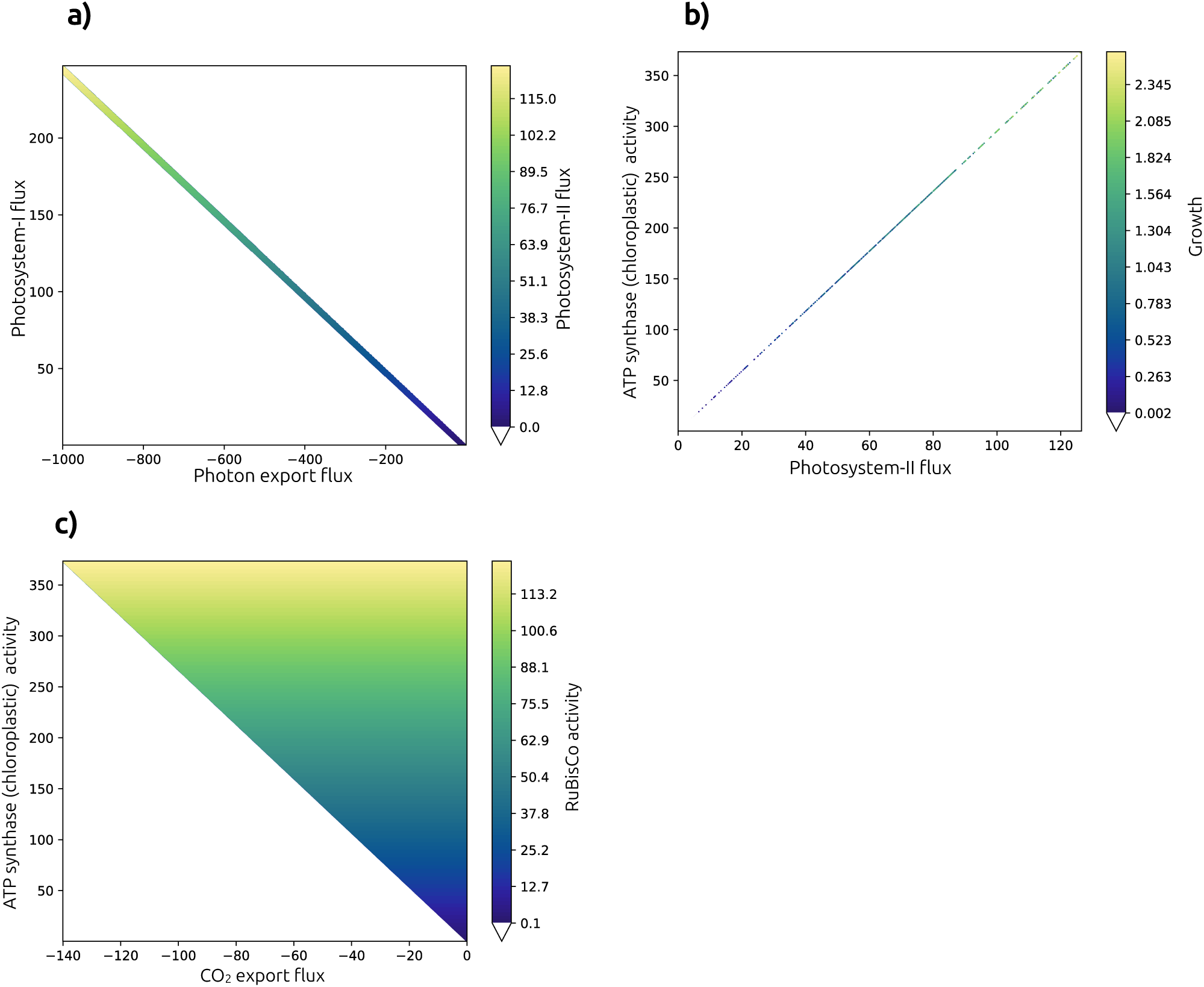

### 2 - Definition of PhotoEukStein-derived GSMs quality

To ensure the biological reliability of the generated models produced by PhotoEukStein, we made sure to classify them into 4 categories. We based our classification on the genome/transcriptome completion, and the frequency of carved reactions (reactions imported without direct genetic evidence). Quality thresholds for PhotoEukStein-derived GSMs is based on the following criteria (cf. Supplementary Table S2).

- Excellent: ≥ 75% completion AND 10% carved reactions
- High: ≥ 50% completion AND < 20% carved reactions
- Medium: ≥ 25% completion AND < 30% carved reactions
- Low: the rest

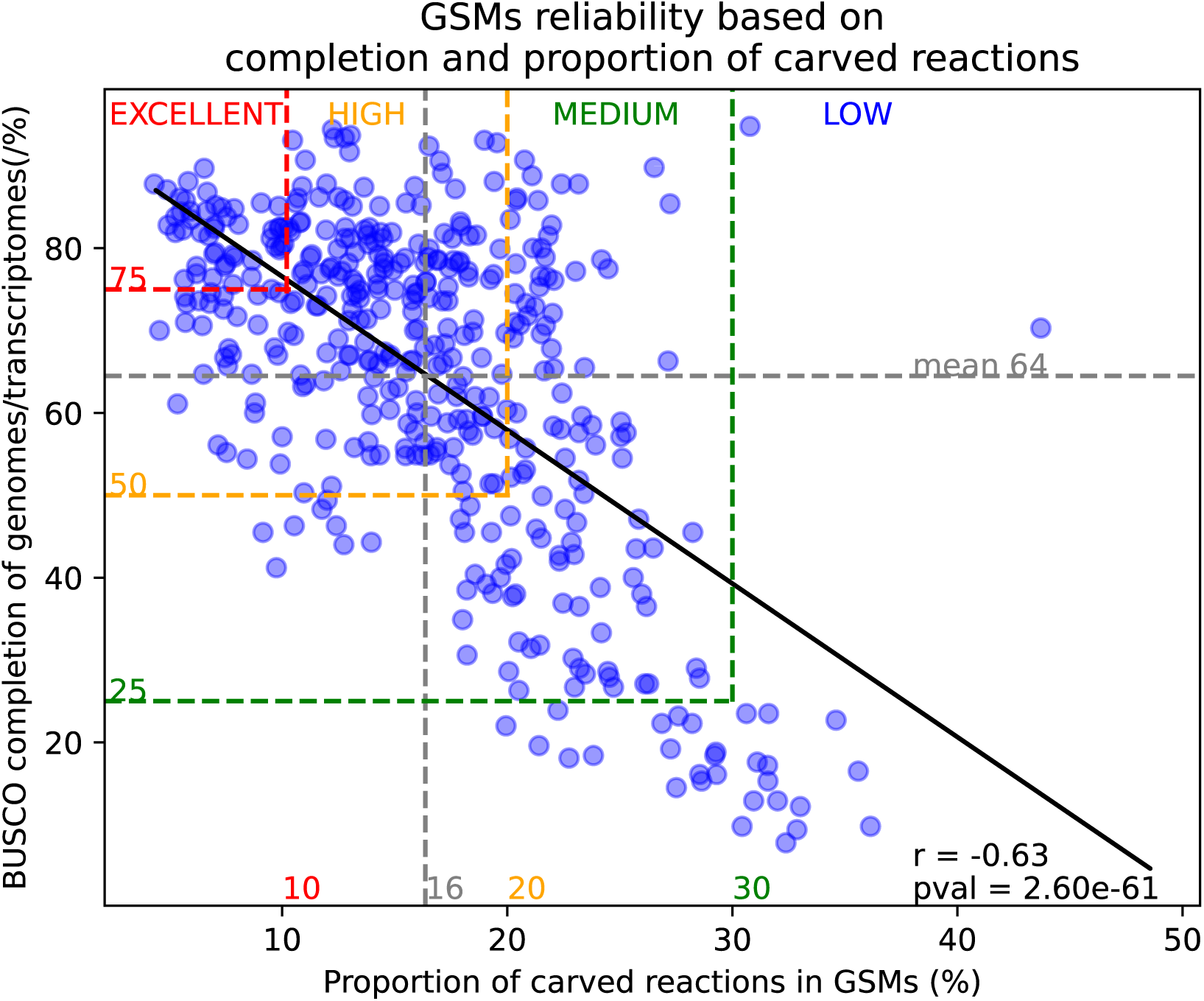

The following figure shows the distribution of PhotoEukStein-based GSMs quality

---

**Supplementary Figure S1.**
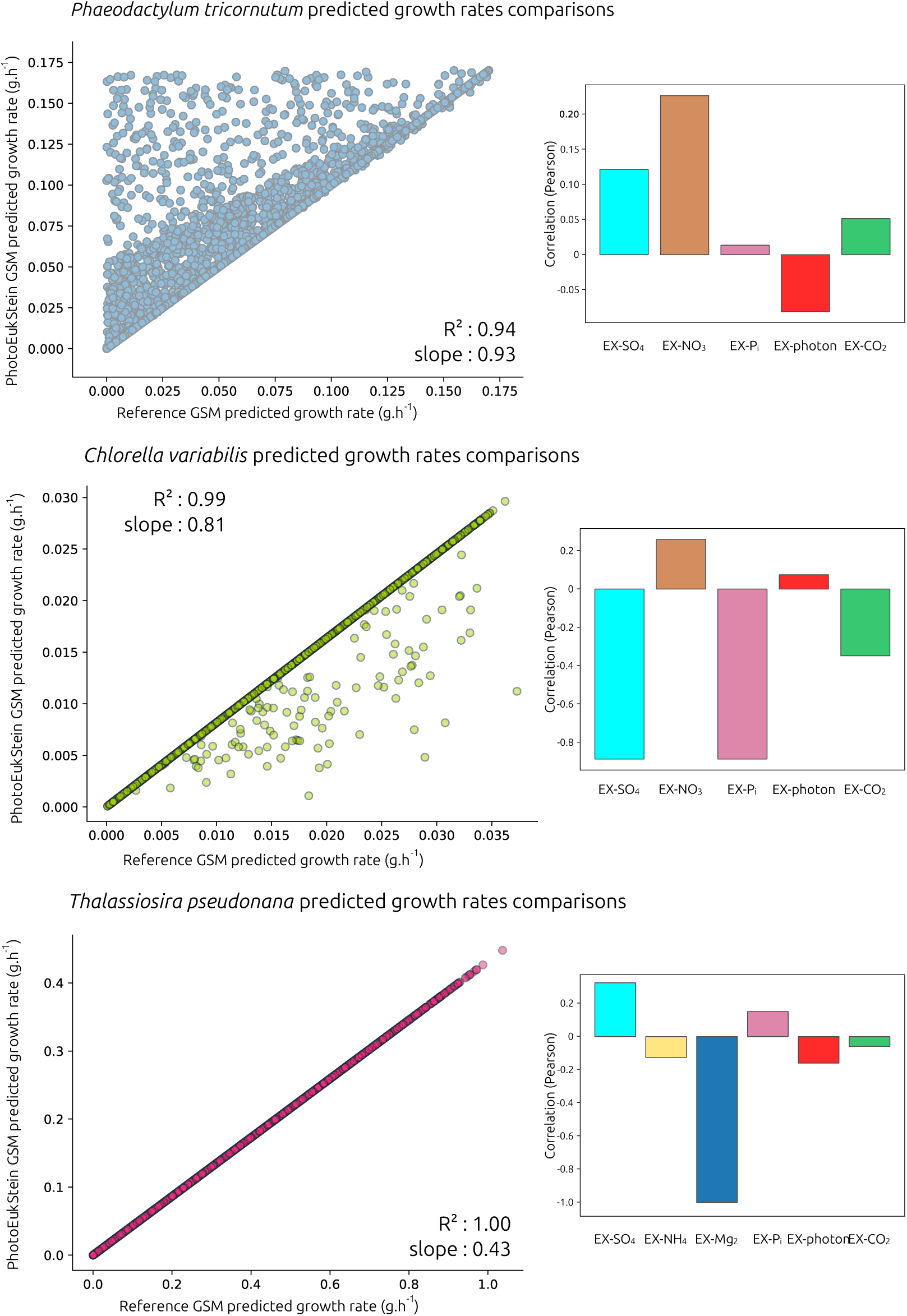
: Comparison of predicted growth rates from PhotoEukStein or reference GSMs for Phaeodactylum tricornutum (top left), Chlorella variabilis (middle left) and Thalassiosira pseudonana (bottom left). In each case, 10,000 iterations of random sampling within each GSMs’ niche space was performed and growth rates predicted for both models and reported. On the right are indicated the metabolites exchange reaction the most correlated with predicted growth rates differences between reference and PhotoEukStein GSMs.

**Supplementary Figure S2.**
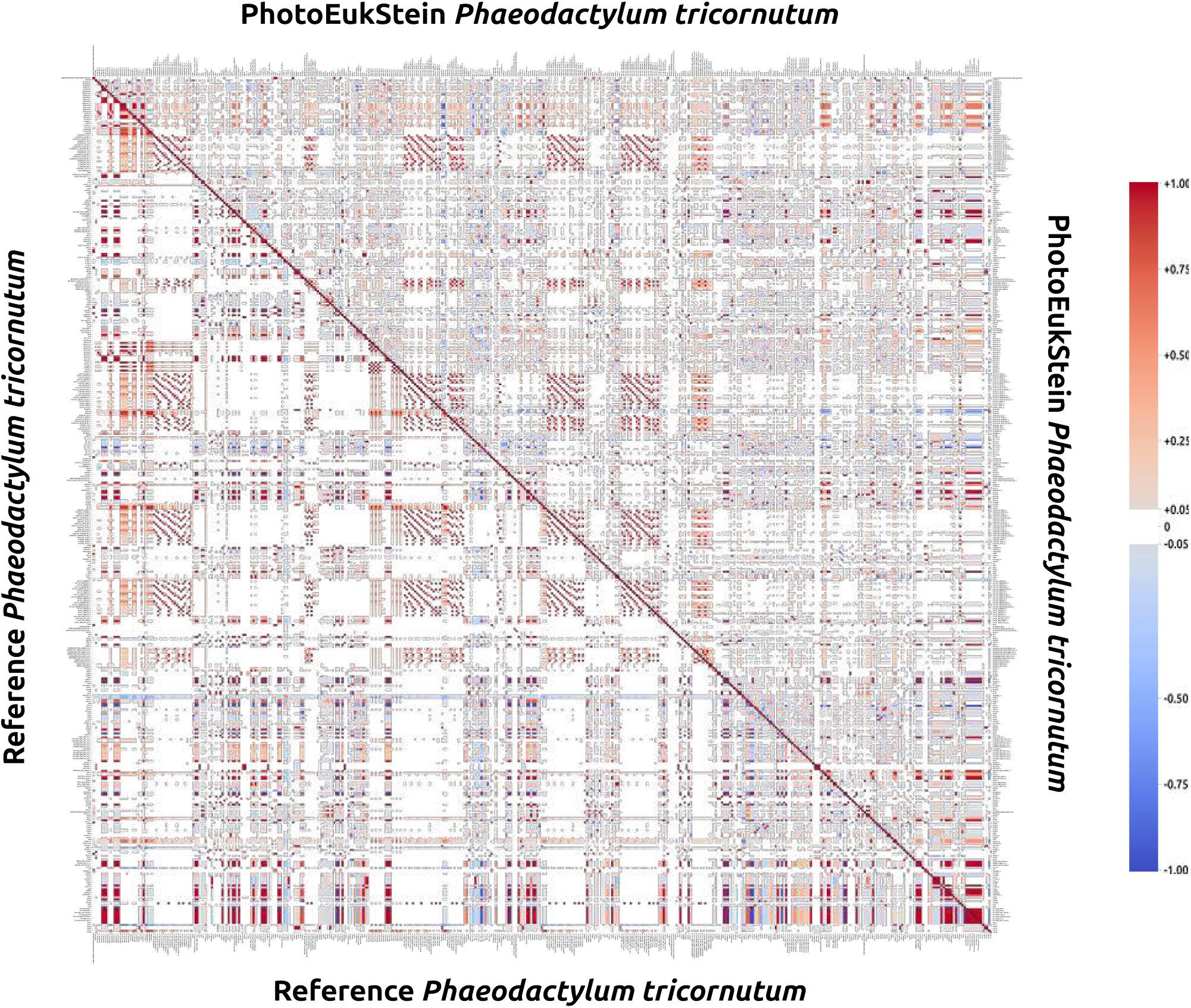
: Correlation matrix comparing original (iLB1034) and PhotoEukStein-derived GSMs reaction fluxes of Phaeodactylum tricornutum. The 434 shared reaction showing at least 1% or their correlations with other reactions fluxes greater to 0.2 when randomly sampling the whole metabolic space are displayed. Lower left rectangle: iLB1034 reactions, upper right rectangle, PhotoEukStein reactions. Only correlation with absolute value greater than 0.5 are coloured.

**Supplementary Figure S3.**
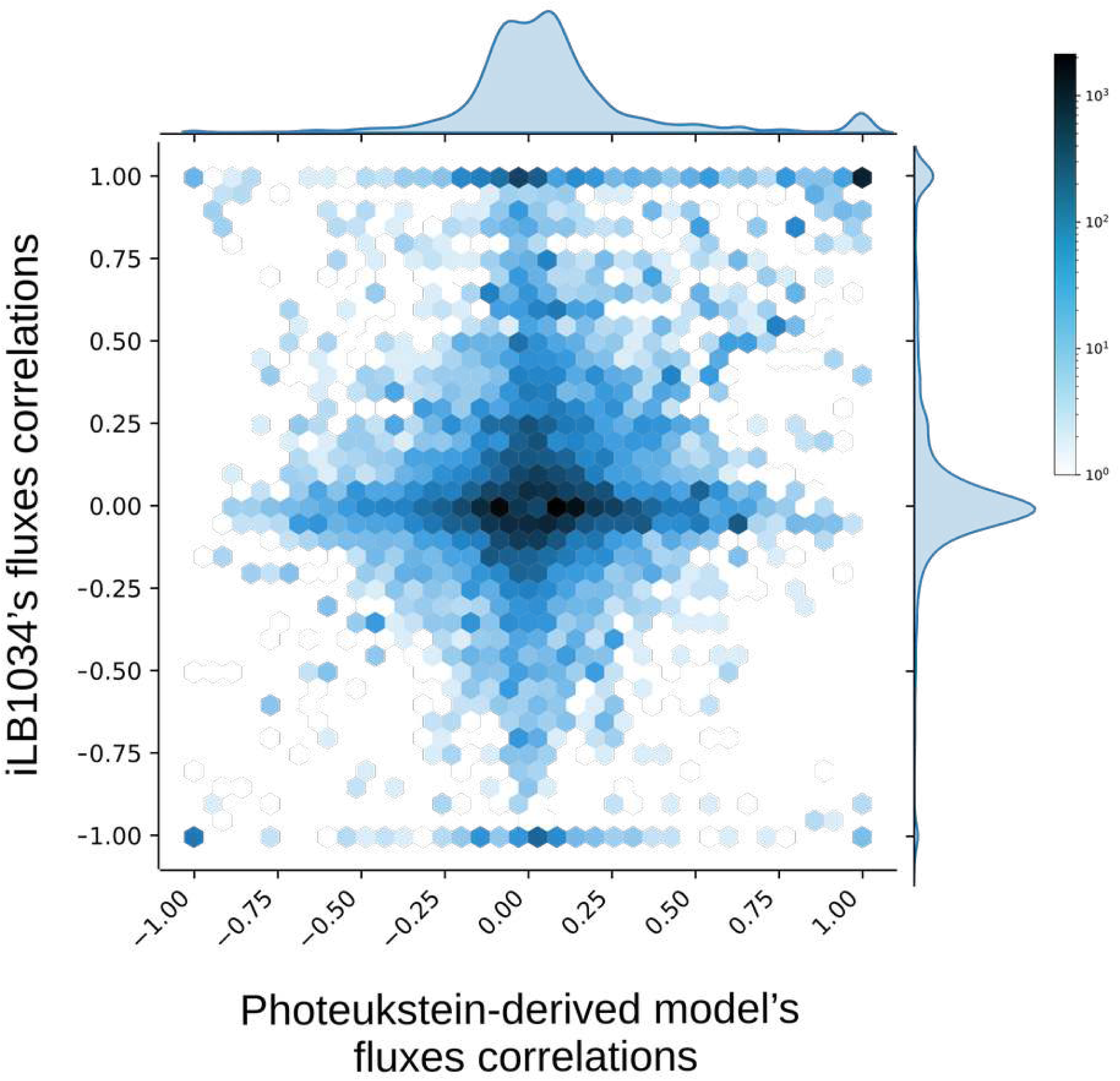
: HexBin representation of distribution of compared reaction fluxes correlations between PhotoEukStein and iLB1034 reference GSMs for Phaeodactylum tricornutum. Effective of correlations pairs of the 434 common reactions showing a possible correlation displayed in Figure S2 (see Materials and Methods section 3). Each cell indicates the number of reactions with corresponding correlations values pairs in iLB1034 (x-axis) and PhotoEukStein (y-axis) during random sampling of metabolic niche space.

**Supplementary Figure S4.**
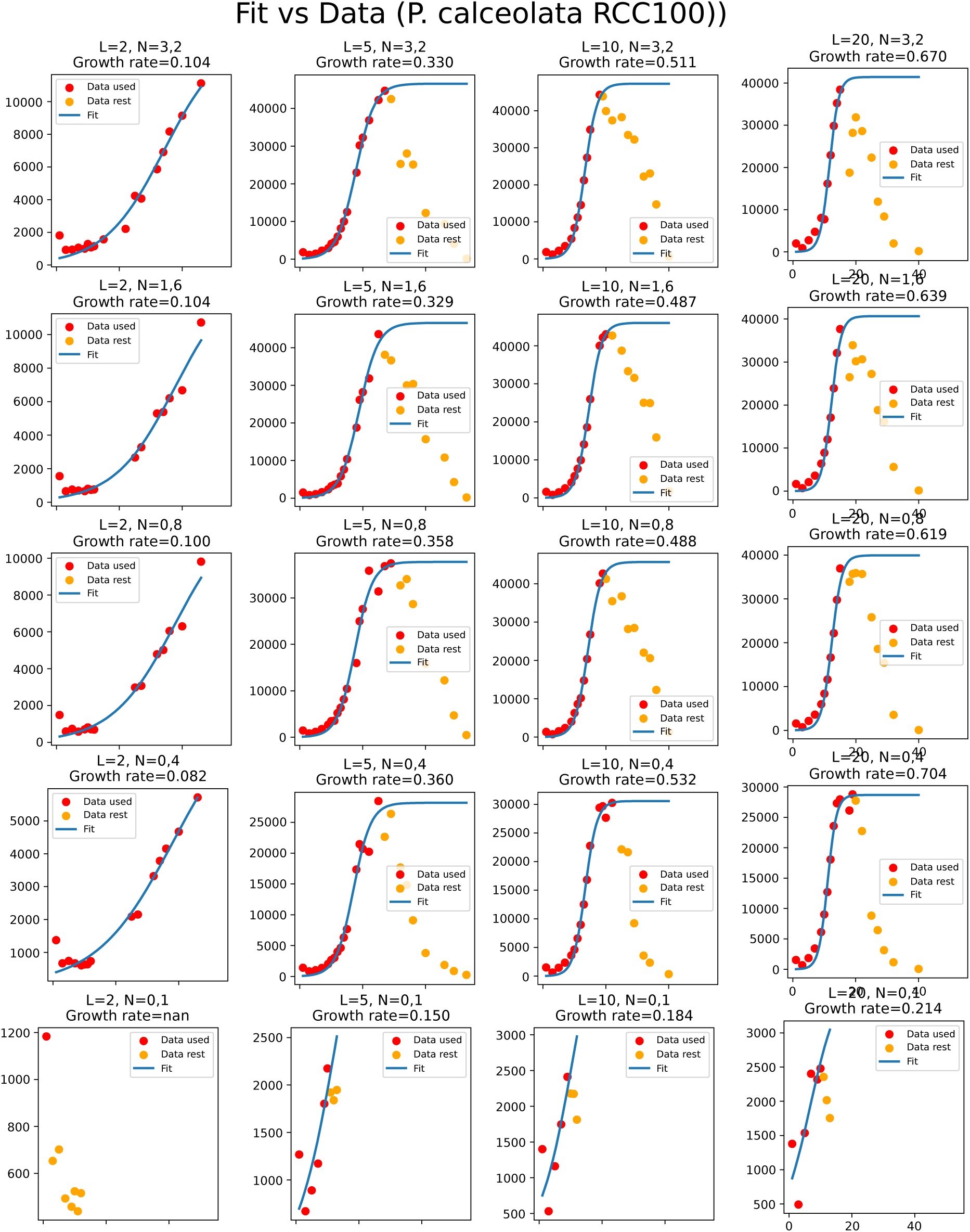
: Experimental determination of growth rates for P. calceolata RCC100

**Supplementary Figure S5.**
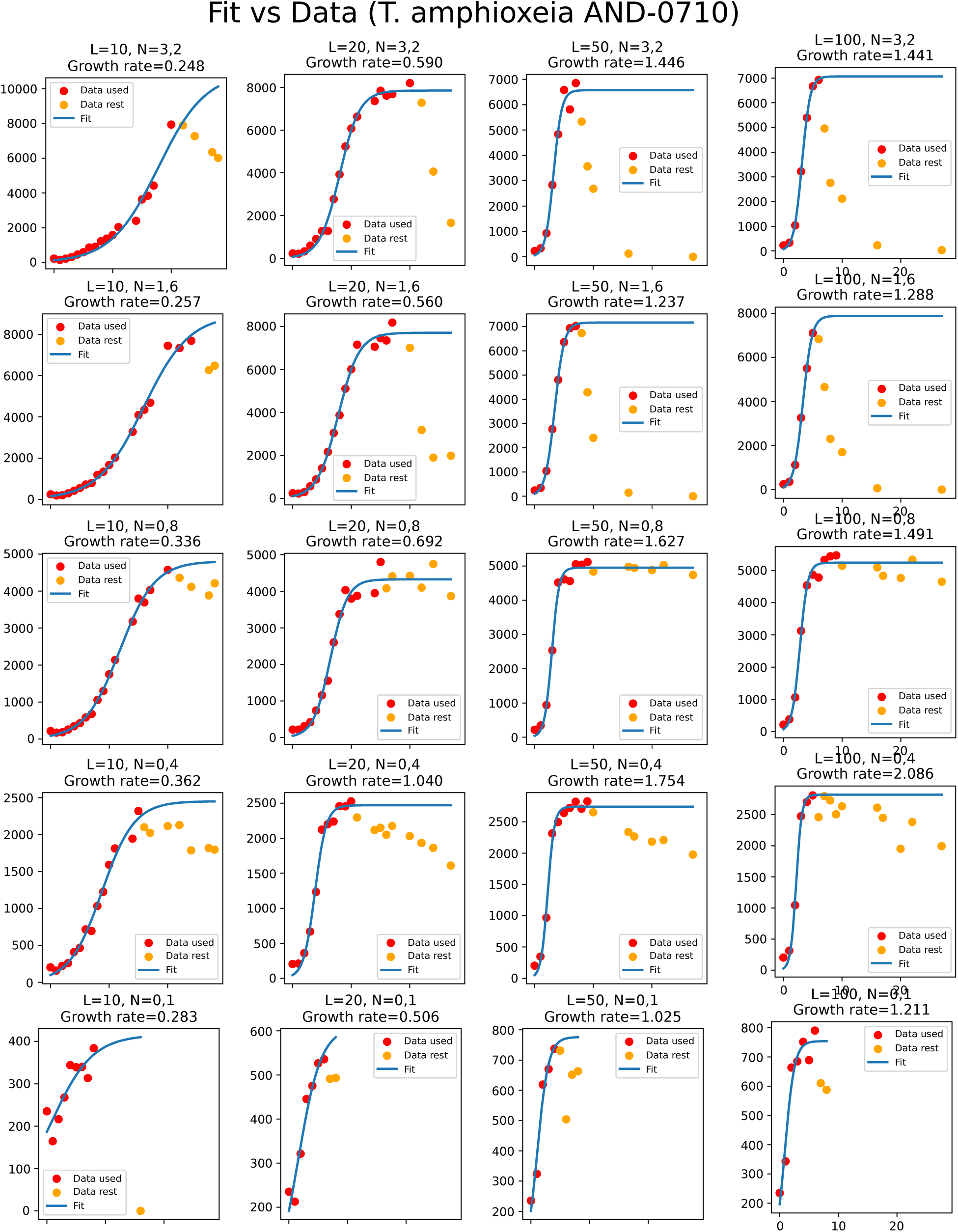
: Experimental determination of growth rates for T. amphioxeia AND-0710

**Supplementary Figure S6.**
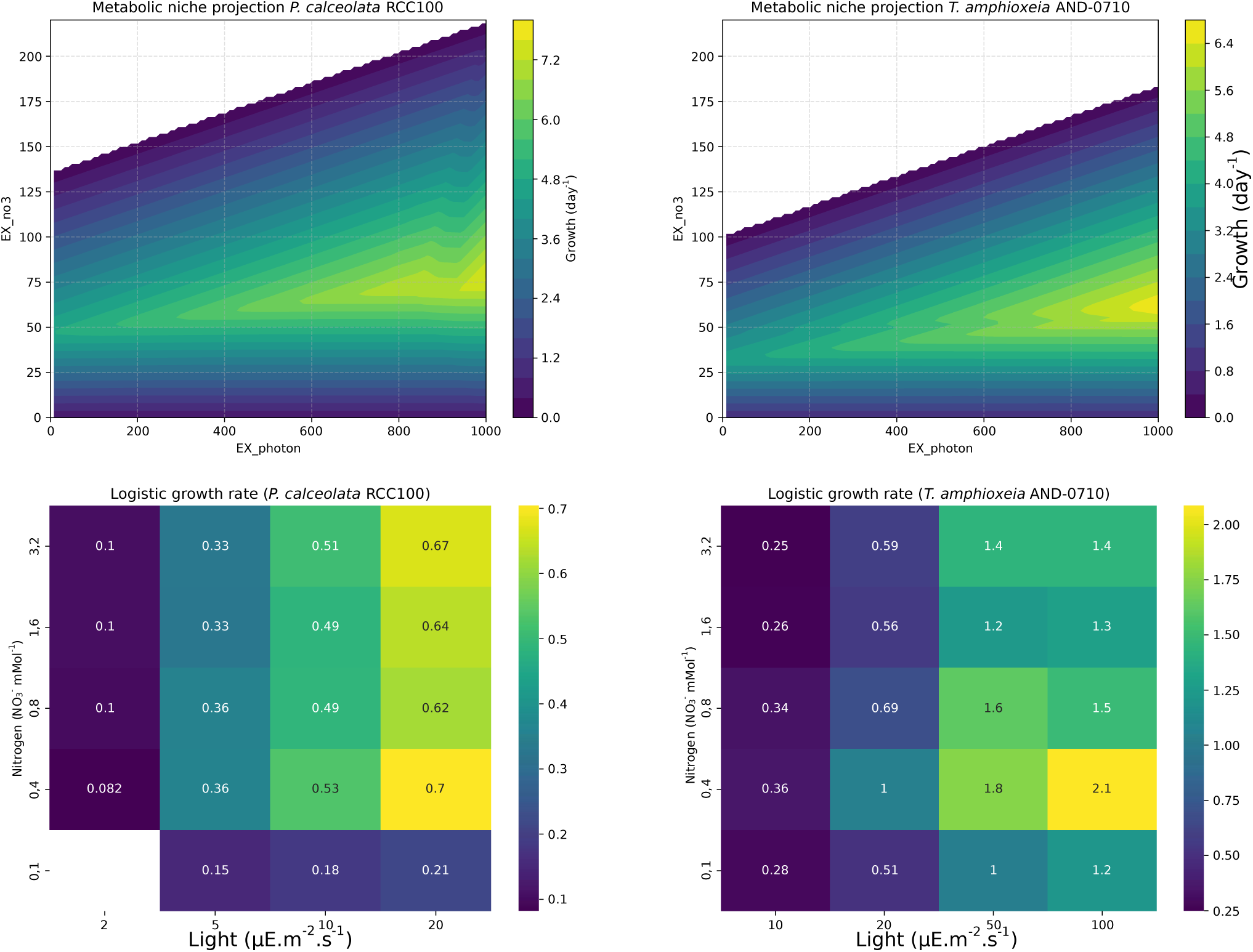
: Comparison of experimentally observed and model-predicted growth rates for Teleaulax amphioxeia AND-0710 and Pelagomonas calceolata RCC100. Heatmaps of logistic growth rate r metabolic niche projections predicted by PhotoEukStein (top) and estimated from culture experiments (bottom) for Pelagomonas calceolata RCC100 (left) and Teleaulax amphioxeia AND-0710 (right across combinations of light intensity and nitrate concentration.

**Supplementary Figure S7.**
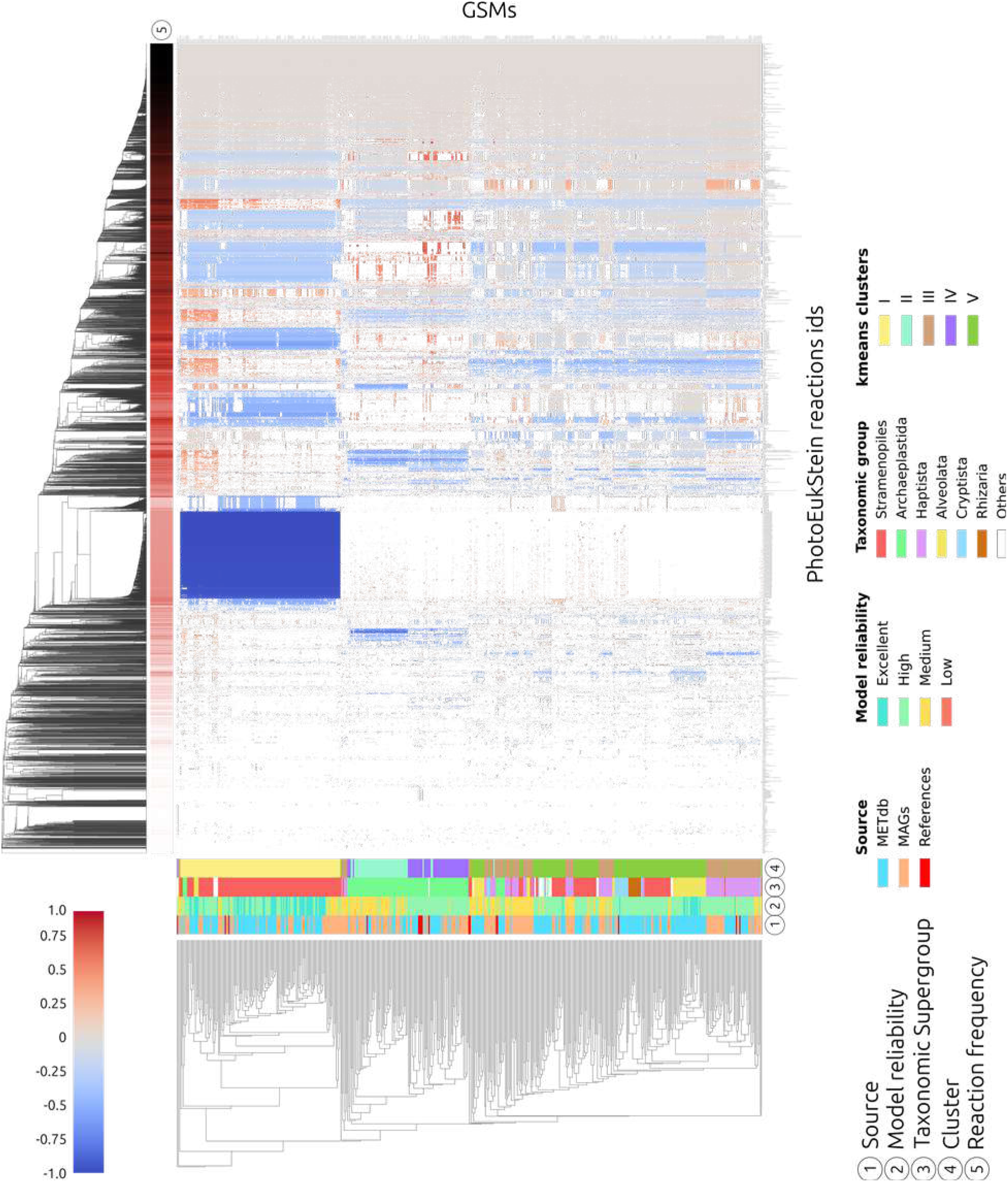
: Compositional analysis of 512 PhotoEukStein- derived GSMs. Presence/absence of 9648 reactions were used to hierarchically cluster (euclidian distance, ward distance) both GSMs (lines) and reactions (columns). Genome source, Taxonomic group, model quality score (as defined in Extended Data), and metabolic cluster (as defined in Figure S5) are shown for each GSM. Frequency of reaction appearance among the 512 GSMs (bad quality models excluded) is indicated for each reaction (from white= 1 to dark red=512). Blue indicates significantly present reactions in a cluster, red if the absence of the reaction is signature reaction, grey if present but not significant.

**Supplementary Figure S8.**
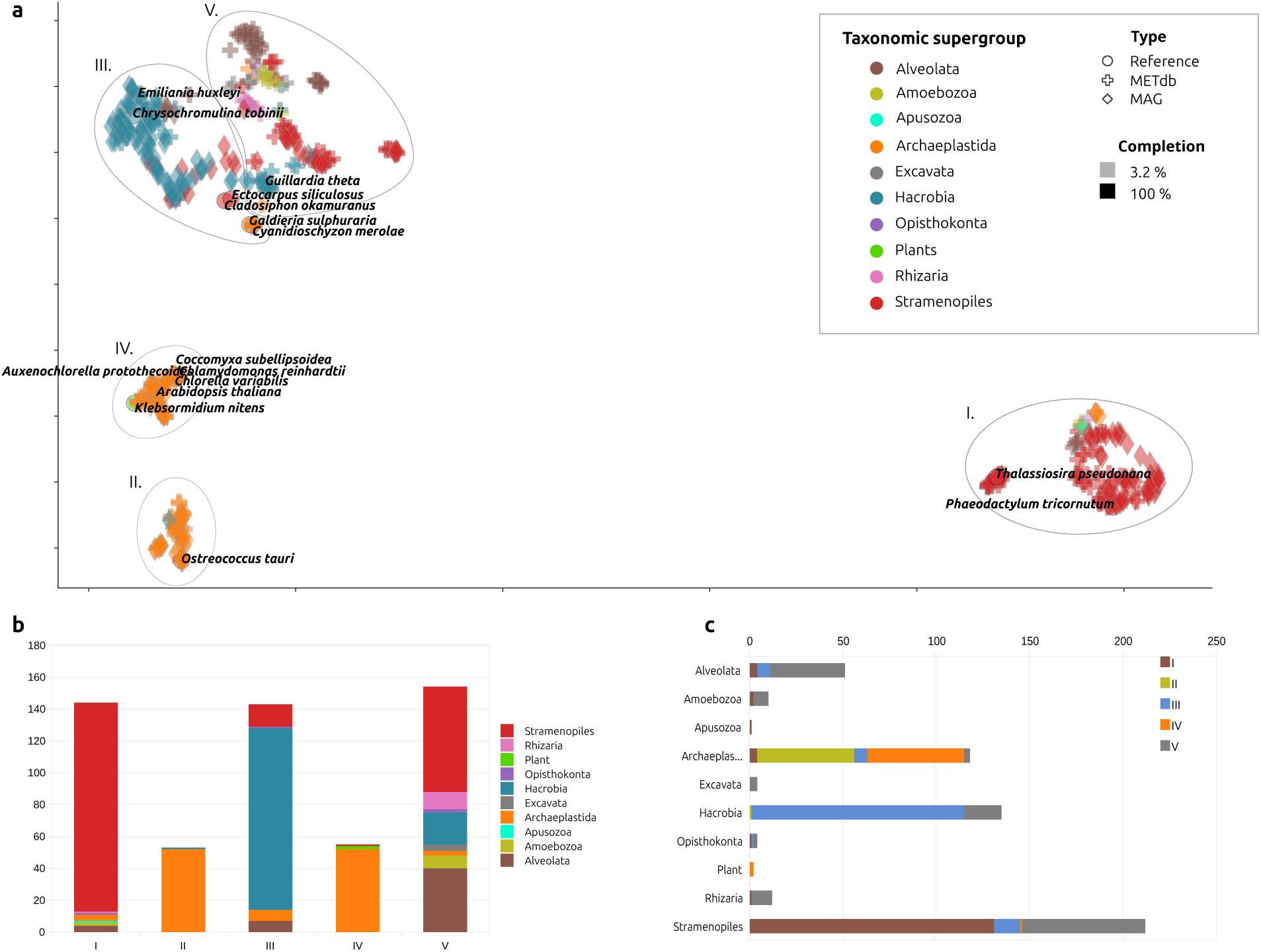
: Compositional analysis of the 549 PhotoEukStein derived GSMs. a) Diffusion map multiscale geometric analysis of GSMs topology followed by umap reduction of dimension processing have been performed to study the distribution of GSMs topologies in the functional space. Shapes indicate origin of the supporting genome (diamonds for MAGs, crosses for METdb and circles for references). Colours indicate main taxonomical groups, and transparency reflect genomes/transcriptomes completions estimations. Grey ellipses identify the 5 clusters supported by k-means signal deconvolution analysis (see Extended Data). b) Repartition of main taxonomic groups within each cluster. c) Distribution of each taxonomic group across the clusters.

## Notes

https://www.genoscope.cns.fr/PhotoEukStein/

### Competing Interest Statement

The authors have declared no competing interest.

### Summary of Updates

We have added an experimental validation step to assess the model predictions on model organisms not included in the building of the system, to better assess the versatility of our system.We explored the impact of nitrate and light depletion on a Pelagophyte (Pelagomonas calceolata RCC100) and a Cryptophyte (Teleaulax amphioxeia AND-A0710). This addition is presented in the newly included Supplementary Figures S4, S5 and S6, as well as Supplementary Tables S11 and S12. We have also expanded the Results and the Discussion sections to reflect these results.

